# Intact cells selectively amplify and structurally remodel amyloid fibrils

**DOI:** 10.1101/2024.09.09.612142

**Authors:** Shoyab Ansari, Dominique Lagasca, Rania Dumarieh, Yiling Xiao, Sakshi Krishna, Yang Li, Kendra K. Frederick

## Abstract

Amyloid forms of α-synuclein adopt different conformations depending on environmental conditions. Although advances in structural biology have accelerated fibril characterization, it remains unclear which conformations predominate in biological settings because current approaches typically require fibrils to be isolated from their native environments. In addition, these approaches provide limited information about flanking flexible regions. Here, using a defined polymorphic seed population and quantitative in-cell NMR, we show that propagation in intact cellular environments—but not in vitro buffer or crowded cellular lysates—reshapes the conformational ensemble of α-synuclein fibrils. In vitro and in cellular lysates, both amyloid-core structure and the conformational preferences of flanking intrinsically disordered regions of the seed are faithfully propagated. In contrast, propagation inside intact cells selectively amplifies the amyloid-core conformer that is minor in vitro, increases its molecular order, and remodels conformational preferences in the flanking disordered region. These results demonstrate that cellular organization plays a decisive role in determining amyloid structure and that biologically relevant conformations cannot be inferred from purified or lysate systems alone.

## Introduction

Neurodegeneration in Parkinson’s disease is associated with Lewy bodies, intracellular inclusions that contain fibrillar aggregates of α-synuclein (α-syn). Although α-syn is an intrinsically disordered monomer in solution, this 140-amino-acid protein can spontaneously assemble into multiple self-templating fibrillar forms that differ in the number and arrangement of β-sheets (*1, 2*). Because α-syn conformations depend strongly on environment, determining which conformations are present in health and disease is critical for understanding pathogenesis and for developing diagnostic and therapeutic strategies. Advances in structural biology have rapidly expanded the number of solved amyloid structures (*2, 3*), raising the possibility of structure-informed intervention. Realizing this promise, however, requires knowing which structures are adopted in biological settings and in what relative abundance (*4*).

A central unresolved question is whether structures determined outside cells accurately represent those present in biological contexts. Current approaches generally rely either on isolating fibrils from biological material or on amplifying fibrils in vitro, but both strategies can distort the underlying conformational ensemble. Amyloid fibrils can be purified from post-mortem tissues based on their high stability and molecular weight (*5, 6*), but these procedures may alter fibril conformations and preferentially enrich the most thermodynamically stable species rather than the most prevalent ones in vivo. Alternatively, pre-formed fibrils can be used to seed amyloid propagation in vitro and in cellular and murine model systems (*7–11*), but the amplified structures need not be identical to the seeds (*10, 12, 13*). Thus, although many amyloid structures have now been characterized, the degree to which these models capture the dominant conformations present in cells remains unclear.

This challenge extends beyond the amyloid core. Nearly half of the residues in α-syn lie outside the ordered core and form flanking intrinsically disordered regions that are poorly resolved by diffraction-based methods and are often described as a “fuzzy coat” surrounding the fibril (*14, 15*). Yet these regions are functionally important: they contribute to primary and secondary nucleation (*16–18*), mediate interactions with cellular components such as chaperones (*19–23*), and influence aggregation, toxicity, and cellular dysfunction (*24, 25*). Understanding amyloid structure in cells therefore requires not only identifying which core polymorphs are present but also determining the conformational preferences of flanking disordered regions that are largely inaccessible to conventional structural approaches.

Magic-angle-spinning (MAS) NMR of frozen solids can provide quantitative information about conformational ensembles because the line shapes of frozen protein samples report on backbone conformational distributions (*26–31*). Unlike most structural methods, NMR does not require purified samples and can therefore be applied in native-like settings to both ordered and disordered regions. Although solid-state NMR is often limited by sensitivity, dynamic nuclear polarization (DNP) dramatically increases signal intensity and enables specific detection of isotopically enriched proteins at biologically relevant concentrations and in complex environments (*29, 32–37*). Our group has established sample handling methods for DNP-assisted MAS NMR that support high-sensitivity measurements on frozen cellular samples that remain intact and viable throughout the experiment (*36, 38–40*). DNP-enhanced MAS NMR can therefore provide quantitative, site-specific information about the full conformational ensemble of a protein both in vitro and inside intact cells.

Here, we directly test whether amyloid conformations are faithfully propagated across environments by starting from a defined seed ensemble containing major and minor fibril polymorphs and quantitatively tracking its evolution. To do so, we use acetylated full-length α-syn(A53T) fibrils (*41, 42*), chosen to match the amino-terminally acetylated α-syn produced in the biosensor cells (*43*), to seed amyloid propagation in buffer, in concentrated cellular lysates, and inside living cells. For cellular propagation, we use a HEK293T biosensor line expressing α-syn(A53T)-CFP/YFP (*43–47*), into which isotopically labeled monomeric α-syn is introduced by electroporation before fibril seeding. This system allows us to compare the amyloid core and flanking disordered regions of the seed ensemble with those of fibrils propagated in vitro, in lysates, and inside cells using a combination of DNP-assisted NMR and cryo-EM. We find that in vitro and lysate conditions faithfully preserve the seed ensemble, whereas propagation inside intact cells selectively amplifies the amyloid-core polymorph that is minor in vitro and remodels the flanking disordered region.

## Results

### Ac-α-syn(A53T) fibrils are polymorphic

To begin our investigation, we characterized the structure of the in vitro formed amyloid fibrils used as seeds for subsequent experiments. Although prior cryo-EM studies of Ac-α-syn(A53T) fibrils reported a single structure (*41*), we sought to test whether fibrils prepared under these conditions were conformationally homogeneous. To assess the conformation of α-syn fibrils by MAS NMR spectroscopy, we used valine chemical shifts as reporters of Ac-α-syn(A53T) amyloid fibril conformation. Following a published protocol (*41*), we formed *de novo* amyloid fibrils using purified recombinant protein that was isotopically enriched with ^13^C at all valines and ^15^N at the single histidine (H50) in the α-syn sequence (**Figure 1E**). As previously reported, the resulting fibrils had a single apparent morphology with helical twist periodicity of 95 ± 2 nm by negative stain transmission electron microscopy (**Figure S1**), consistent with a structurally uniform preparation.

**Figure 1:**
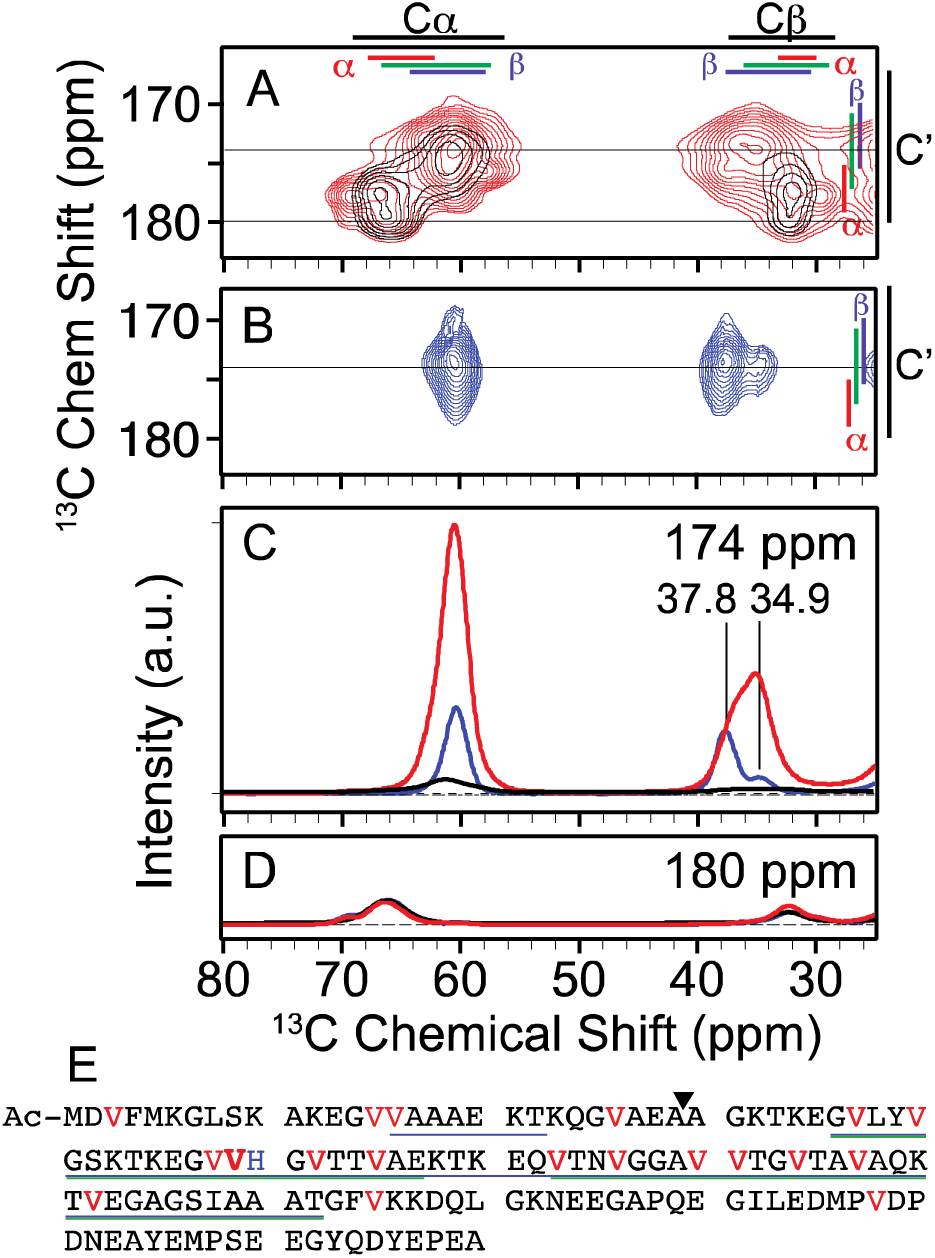
A) Overlay of 2D ^13^C–^13^C DARR spectra of uniformly ^13^C-valine labeled Ac-α-syn(A53T) fibrils (red) and segmentally ^13^C-valine labeled fibrils where only the four N terminal valines (black) are labeled. Average chemical shifts +/- two standard deviation are marked for each atom (black) as well as for the α-helical (red), random coil (green), and β (blue) secondary structural elements. B) NCOCx spectra of uniformly ^13^C-valine and ^15^N-histidine labeled fibrils that is specific for V49 (blue). One-dimensional slice from the 2D spectra at C) 174 ppm and D) 180 ppm. The intensity of the V49 spectrum (blue) is displayed at 10 times the relative intensity for visibility and the peak centers are indicated. NMR spectra were recorded at a field strength of 14 T (600 MHz) and MAS frequencies of 12 kHz at 104 K. NMR samples were cryoprotected with 15% d_8_-^12^C-glycerol and doped with 7 mM AMUPol. E) Primary sequence of Ac-α-syn(A53T). The 19 valines (highlighted in red) are distributed throughout the sequence, with V49 bolded for emphasis. H50 is colored blue. Regions visible in the cryo-EM density for polymorph A (blue) and polymorph B (green) are underlined. The ligation site for segmental isotopic labeling is indicated by a triangle.

To assess conformational homogeneity at a single site in the amyloid fibril, we collected 2D NCOCx spectra at 104 K (*48*) (**Figure 1B**). This experiment is selective for V49 given the labeling pattern described above. The chemical shifts of V49 are consistent with β-sheet conformations (*49*) and revealed two distinct conformations at this site, with populations of 79% and 21% for the major and minor polymorphs, respectively (**Figure 1B**; **Table S1**). Predictions of the φ/ψ backbone torsion angles of V49 from chemical shift and homology (*50*) indicated that the conformational differences between the two polymorphs are small; the predicted dihedral angles at V49 were -132°/138° ± 20° for the major polymorph and -113°/136° ± 20° for the minor polymorph, a change that displaces the adjacent residues by less than 1 Å (**Figure 2C**). The major peak (V49Cβ) had a full width at half maximum (FWHM) of 2.0 ppm, which is ∼0.5 ppm broader than the expected homogeneous linewidth under these conditions, and the minor peak (V49Cβ’) had a FWHM of 3.1 ppm (**Table S1**). The broader linewidth indicates that the minor conformation samples a wider range of related conformations than the major conformation (*26–28, 51*). Thus, site-specific NMR reveals that an apparently homogeneous fibril preparation is in fact composed of multiple conformations and enables quantitative determination of the relative populations of these conformations with atomic-level precision.

**Figure 2:**
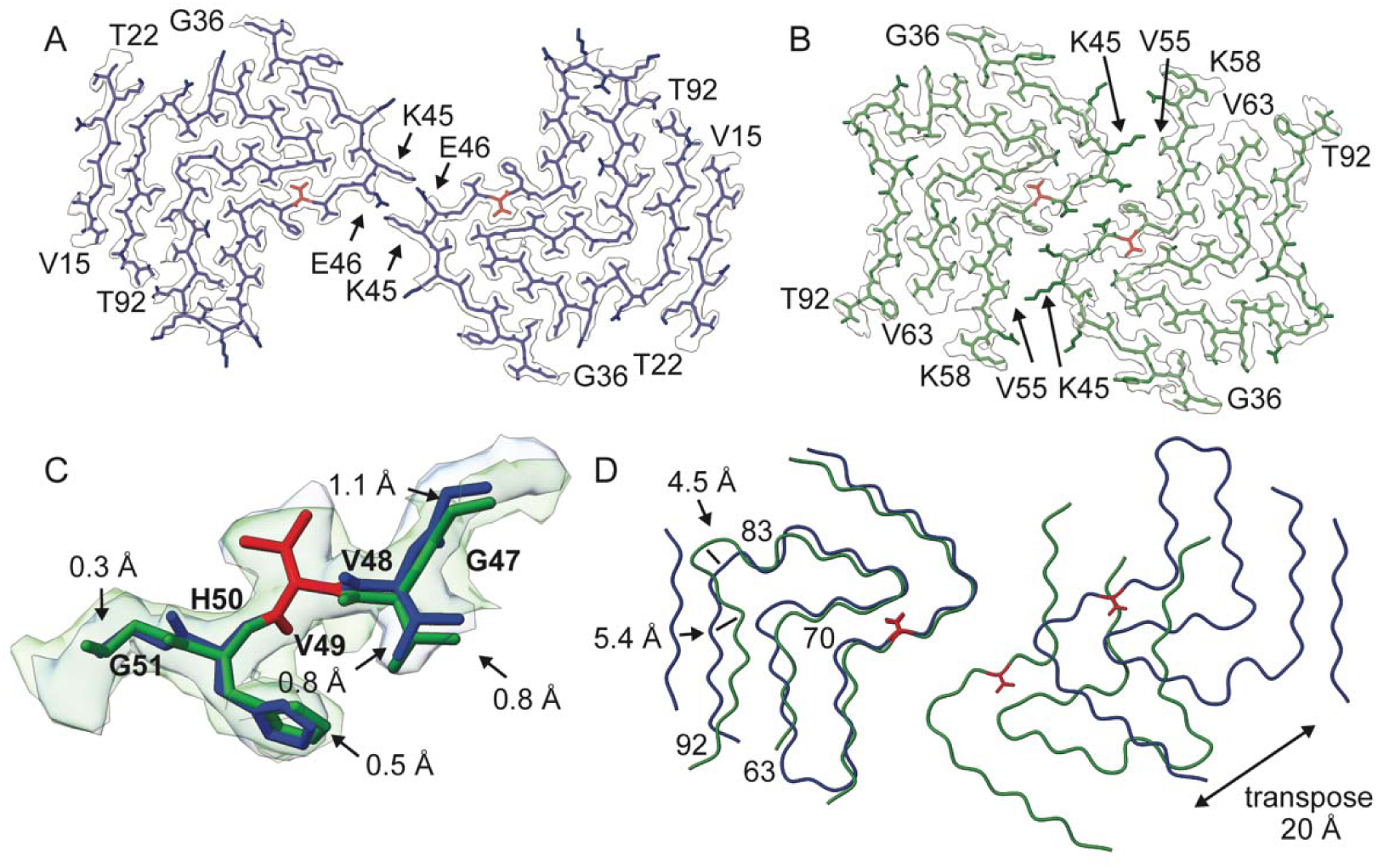
**Cryo-EM analysis of Ac-**α**-syn(A53T) fibrils reveals two distinct polymorphs.** One layer of the fibril structure model in the cryo-EM reconstruction density map of A) polymorph A (blue) and B) polymorph B (green) with V49 highlighted in red and the protofilaments interfaces annotated with arrows. C) Local structural variation (G47 to G51) between polymorph A and B when V49 (red) is aligned. D) Overlay of a single layer of polymorph A (blue) and polymorph B (green), based on the alignment of the Cα atoms of one protofilament subunit, highlight small differences in the protofilament fold (e.g. residues 63-70 and 83-92) and a large rearrangement of the protofilament interface due to a ∼ 20 Å transposition of the second protofilament. These structural differences correspond to distinct conformational states identified by NMR.

To determine whether the polymorphism observed by NMR was limited to V49 or reflected larger-scale structural differences, we collected single-particle cryo-EM data for the Ac-α-syn(A53T) fibrils. Two polymorphs were distinguished in the 2D class averages, and the reconstructed cryo-EM 3D density maps had overall resolutions of 2.2 Å for polymorph A and 2.7 Å for polymorph B (**Figure 2, Figure S2, Table S2**). The polymorphism observed by cryo-EM was not limited to V49; these two polymorphs differed in the dihedral angles at V49 — the φ/ψ backbone torsion angles for V49 in polymorph A were -131°/134° and in polymorph B were -106°/136° (**Figure 2C**) — as well as in the number of residues sequestered in the amyloid core (14–22 & 36–92 vs. 36–92), the main-chain traces (e.g. residues 63–70 and 83–92), and, most strikingly, the protofilament interfaces (**Figure 2D**). While fibrils of both polymorphs consisted of two protofilaments with a Greek-key-like architecture (r.m.s.d. = 0.8 Å for 68% of the residues), the protofilament interface of polymorph A involved only K45 and E46 and buried ∼139 Å^2^ of solvent-accessible surface area per monomer, whereas the protofilament interface of polymorph B spanned residues 45–55 and buried ∼316 Å^2^. Thus, the polymorphism detected at V49 reports on both local and large-scale rearrangements of fibril structure, including substantial differences in protofilament interfaces and solvent-exposed surfaces.

Site-specific NMR measurements provide quantitative information about conformational ensembles. NMR and cryo-EM provide complementary information that links population-weighted observables to full atomistic structures. We establish this connection using two independent NMR observables: chemical shifts, which report on backbone dihedral angles, and linewidths, which report on molecular order and can be directly compared to cryo-EM reconstruction behavior. Because the labeling scheme isolates V49, the corresponding chemical shifts can be directly compared with the local backbone geometry of the two cryo-EM polymorphs. The predicted dihedral angles from chemical shifts for the major polymorph agreed with those for polymorph A, while those for the minor polymorph agreed with polymorph B, allowing assignment of the two conformations to the corresponding cryo-EM structures. Moreover, the narrower NMR peak width for the major polymorph indicates higher molecular order, whereas the broader peak width for the minor polymorph reflects greater conformational heterogeneity. Consistent with this, a substantially larger fraction of particles assigned to polymorph A could be used for high-resolution 3D reconstruction (∼10%) than polymorph B (∼1%) (**Figure 2, Table S2**), indicating that polymorph A is more conformationally uniform while polymorph B is more heterogeneous.

Having established the correspondence between NMR signals and cryo-EM structures, the relative intensities of the NMR peaks can be used to determine the populations of the two polymorphs. Because the NMR measurements were performed on frozen samples under the same cryogenic conditions used for cryo-EM, the spectra report directly on the full conformational ensemble present in the sample. NMR measurements therefore established that polymorph A constituted the majority of the conformational ensemble (∼79%), whereas polymorph B represented a minor population (∼21%). This distribution is not reflected in the cryo-EM 2D class particle counts: polymorph B accounts for a larger fraction of particles in the 2D classes (∼55%) than polymorph A (∼45%) (**Table S2**). This discrepancy shows that the number of particles assigned to each polymorph in cryo-EM analyses does not directly reflect their relative abundance in the sample; particle counts reflect not only abundance but also biases introduced during grid preparation and particle selection. Thus, polymorph A represents the major conformation observed by NMR spectroscopy for *de novo* Ac-α-syn(A53T) fibrils *in vitro* while polymorph B represents the minor conformation. Together, these data establish that quantitative spectroscopic measurements are required to accurately determine the relative populations of structural polymorphs and that apparently uniform fibril preparations can conceal substantial structural heterogeneity.

### Flanking disordered regions of amyloid fibrils have conformational preferences *in vitro*

Having established polymorphism in the amyloid core, we next asked whether the flanking regions of the fibrils, which are not resolved by cryo-EM, also exhibit measurable conformational preferences. Of the 19 valines in α-syn, approximately three-quarters are located in the β-sheet- and turn-rich amyloid core, whereas the remaining quarter are in the flanking regions that are unresolved in the cryo-EM structures. To assess the conformations sampled by the valines in our amyloid fibrils, we collected a 2D ^13^C–^13^C dipolar assisted rotational resonance (DARR) correlation spectrum at 104 K. The majority of the cross peak intensity (82% ± 3%) had chemical shifts consistent with β-strand and random coil dihedral angles (*49*) **(Figure 1**, red). However, a minority of the cross peak intensity (18% ± 3%) had chemical shifts consistent with α-helical dihedral angles (*49*). Because the valines in the amyloid core are in β-strands and turns, these data suggested that the valine residues in the amino terminal region may adopt dihedral angles found in α-helical conformations.

To determine if the flanking disordered region of Ac-α-syn(A53T) fibrils had strong preferences to adopt dihedral angles found in α-helical conformations, we formed amyloid fibrils from Ac-α-syn(A53T) isotopically enriched with ^13^C at only the four valines in the amino terminal region (V3, V15, V16, and V26) and collected a 2D ^13^C-^13^C DARR spectrum at 104 K (**Figure 1**, black). Most of the amino terminal region (residues 1-36) was unresolved in the cryo-EM structure of both polymorphs, although in polymorph A, a short segment (V15-T22) formed a β-strand that encompassed two of the four isotopically labeled valines (V15 & V16) (**Figure 2A**). In the segmentally isotopically labeled sample, most of the cross-peak intensity for the four valines in the amino terminus had chemical shifts consistent with α-helical dihedral angles (67% ± 8%) whereas a minority of the cross-peak intensity had chemical shifts consistent with β-strand and random coil dihedral angles (32% ± 8%) (**Figure 1C, 1D**). Comparison of the spectra of the segmentally isotopically labeled Ac-α-syn(A53T) fibrils with those of uniformly ^13^C valine labeled Ac-α-syn(A53T) fibrils revealed that the majority (∼80%) of the cross-peak intensity in the α-helical region of the 2D DARR spectrum of uniformly ^13^C valine labeled fibrils was accounted for by the four amino terminal valine residues (**Figure 1**). Therefore, the peaks in the α-helical region of *de novo* fibrils *in vitro* report on the amino-terminal disordered flanking region. These data show that although this region is unresolved by cryo-EM, it exhibits strong conformational preferences in vitro and is directly observable by DNP-enhanced NMR.

### Ac-α-syn-(A53T) fibrils propagate faithfully *in vitro*

To determine whether the conformational ensemble of de novo Ac-α-syn(A53T) fibrils is faithfully propagated in vitro, we seeded monomeric isotopically labeled Ac-α-syn(A53T) with a small amount of pre-formed de novo fibrils (1% monomer/monomer). The resulting fibrils yielded 1D NCOCx and 2D ^13^C–^13^C DARR spectra that were indistinguishable from those of the de novo formed Ac-α-syn(A53T) fibrils (**Figure S3, Table S4**). These data indicate that in vitro propagation faithfully preserves the full seed ensemble, including both the populations of the amyloid-core polymorphs and the conformational preferences of the amino-terminal flanking region.

### Cellular propagation of Ac-α-syn(A53T) fibrils is efficient and cytotoxic

To determine whether the conformation of Ac-α-syn(A53T) fibrils could be faithfully propagated inside cells, we used Ac-α-syn(A53T) fibrils to seed amyloid formation in a biosensor cell line developed to propagate α-syn amyloid conformations in response to exposure to exogenous seeds (*44*). These HEK293T cells stably express soluble α-syn(A53T) fused through an 18-amino-acid linker to either CFP or YFP (*43, 45–47*), a modification that does not alter seeding efficiency (*42*). Upon aggregation of α-syn(A53T), CFP and YFP are brought into proximity and act as a FRET pair, allowing aggregate formation to be visualized and quantified by fluorescence microscopy and flow cytometry. Seeding of the α-syn(A53T) biosensor cells with Ac-α-syn(A53T) fibrils was efficient: 71.1% ± 0.4% of cells were FRET-positive 24 h after exposure to fibril seeds (**Figure 3A** **& 3B**).

**Figure 3:**
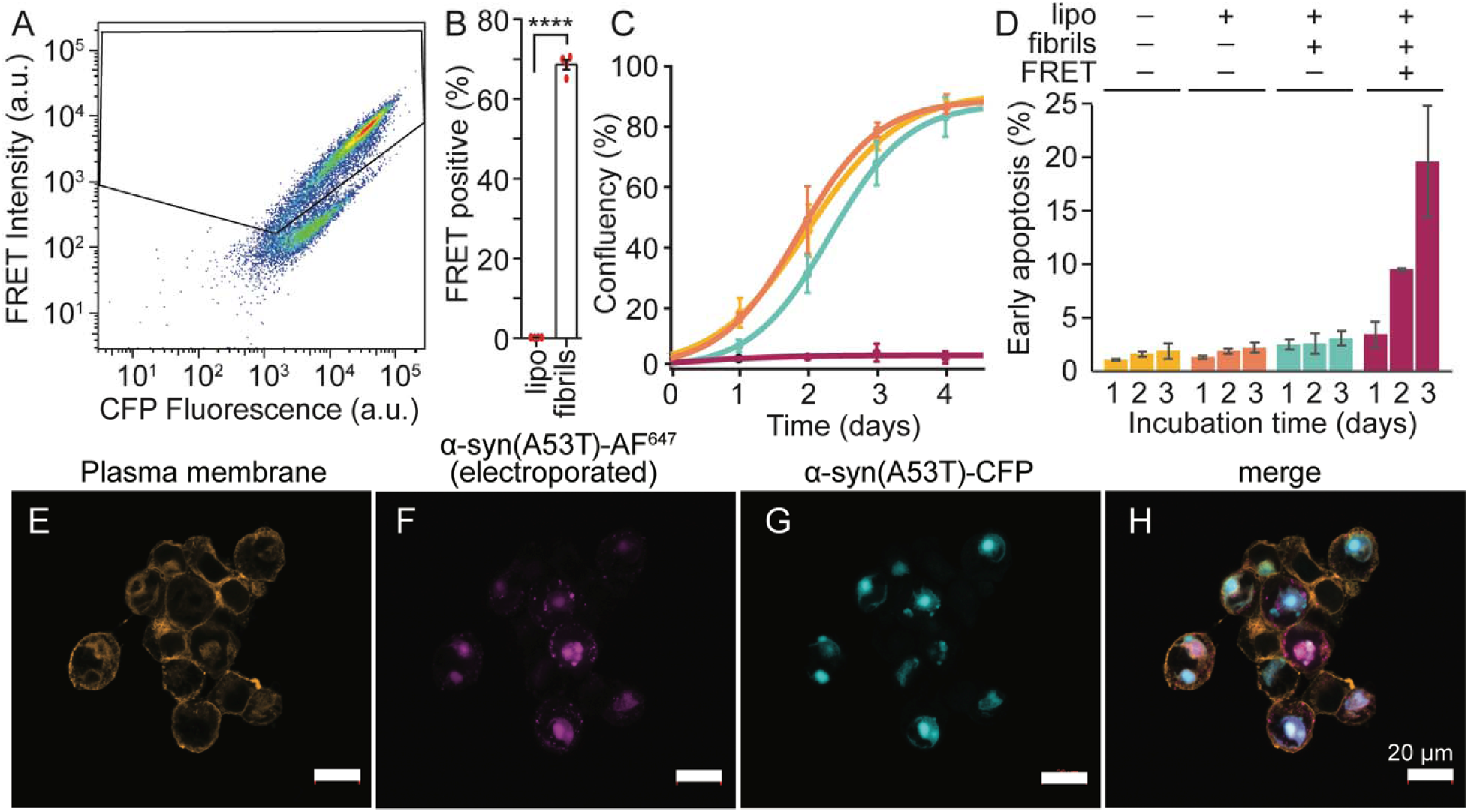
Seeded propagation of Ac-α-syn(A53T) in α-syn biosensor cells is efficient, cytotoxic, and incorporates exogenously delivered protein. A) Flow cytometry reports on seeding efficiency of Ac-α-syn(A53T) preformed fibrils for the α-syn(A53T)-CFP/YFP HEK293T biosensor cell line after 24 hours of incubation. B) Quantification of cytometry experiments, p = 0.0001, paired t-test of fibril-treated versus lipofectamine treated cells. Error bars are ± S.E.M; n=4. C) Cells harboring aggregates do not propagate. Cultured FRET-negative cells after exposure to buffer (yellow), lipofectamine (orange), or lipofectamine and fibrils (blue) grew to confluency after 4 days while FRET-positive cells exposed to lipofectamine and fibrils (pink) did not. Data are average ± standard deviation; n=3. D) The population of cells in early apoptosis as assessed by Annexin V and PI staining increases with increasing incubation time. Data are average ± standard deviation; n=4. E-H) Colocalization of electroporated Alexa 647–tagged Ac-α-syn(A53T) with endogenously expressed Ac-α-syn(A53T)-CFP in biosensor cells after seeding. Exogenously produced Ac-α-syn(A53T) labeled with Alexa fluor 647 (AF^647^) was electroporated into the biosensor cells (pink) and, upon exposure to Ac-α-syn(A53T) fibrils, colocalized with the endogenously expressed Ac-α-syn(A53T)-CFP aggregates (cyan) (Pearson’s coefficient = 0.62).

To determine whether the cellular aggregates formed in response to Ac-α-syn(A53T) seeds were cytotoxic, we sorted fibril-treated biosensor cells into FRET-negative and FRET-positive populations and measured their growth kinetics. FRET-negative cells grew to confluency within 4 days, whereas FRET-positive cells did not (**Figure 3C**), indicating that the presence of aggregated α-syn(A53T)-CFP/YFP suppresses cell growth. We next quantified apoptotic cell death in FRET-negative and FRET-positive cells by annexin V and propidium iodide staining followed by flow cytometry (*52*). Whereas 2.4% ± 0.5% of FRET-negative cells were in early apoptosis 24 h after exposure to fibrils and this fraction remained unchanged over time, 3.4% ± 1.2% of FRET-positive cells were apoptotic at 24 h and this fraction increased to 19% ± 5% after 3 days (**Figure 3D**). Thus, Ac-α-syn(A53T) fibril-induced aggregation in the biosensor cells is both efficient and cytotoxic.

To determine whether exogenously introduced Ac-α-syn(A53T) behaves like the Ac-α-syn(A53T)-CFP/YFP produced by the cells, we introduced recombinant Ac-α-syn(A53T) into the biosensor cells by electroporation using established protein-delivery protocols (*53–55*) delivering 70 ± 4 µM Ac-α-syn(A53T) to the cells (**Figure S5**). Delivery of exogenous Ac-α-syn(A53T) did not alter the aggregation behavior of the biosensor cells: cells remained FRET-negative before exposure to fibrils and became FRET-positive only after fibril addition (**Figure S6**). We then asked whether the exogenously delivered protein was incorporated into the same aggregates as the cell-produced reporter. To do so, we introduced a cysteine mutation at position 122 (*56*), labeled Ac-α-syn(A53T) with Alexa Fluor 647 (AF647), and delivered the labeled protein to cells by electroporation. Following exposure to Ac-α-syn(A53T) fibrils, Ac-α-syn(A53T)-AF647 colocalized with aggregates formed by the endogenously expressed Ac-α-syn(A53T)-CFP/YFP as assessed by confocal microscopy (**Figure 3E-H**), indicating that the exogenously introduced protein enters the same fibril-induced aggregation pathway as the cell-produced reporter.

### Cells selectively propagate and order Ac-α-syn(A53T) polymorph B

To determine which amyloid conformations are propagated inside cells, we delivered exogenously prepared monomeric Ac-α-syn(A53T) specifically enriched with ^13^C valine and ^15^N histidine to the biosensor cells and induced fibril formation by exposing the cells to Ac-α-syn(A53T) seeds. To assess the conformational homogeneity at V49 in the amyloid core, we collected one dimensional NCOCx (*37, 48*) spectra at 104 K on three independently prepared samples. The average signal-to-noise ratio of these spectra was 8:1 at the V49Cα peak and co-added spectrum had a signal-to-noise ratio of 13:1 (**Figure S7**). In all three replicates, as well as in the co-added spectrum, the peak centers of V49Cβ and V49Cβ’ were the same as those of *de novo* fibril seeds, indicating that the same amyloid fibril core conformations were present in both the seed and in the in-cell propagated samples. However, the relative populations of the two conformers were markedly different. (**Figure 4, Figure S7, Table S4, Table S5**). The population of the major peak in the seed population decreased from 79% to 33% whereas the population of the minor peak increased from 21% to 67%. Thus, the minority conformation in the seed population became the majority conformation when amyloid fibrils were propagated inside cells (**Figure 4A**).

**Figure 4:**
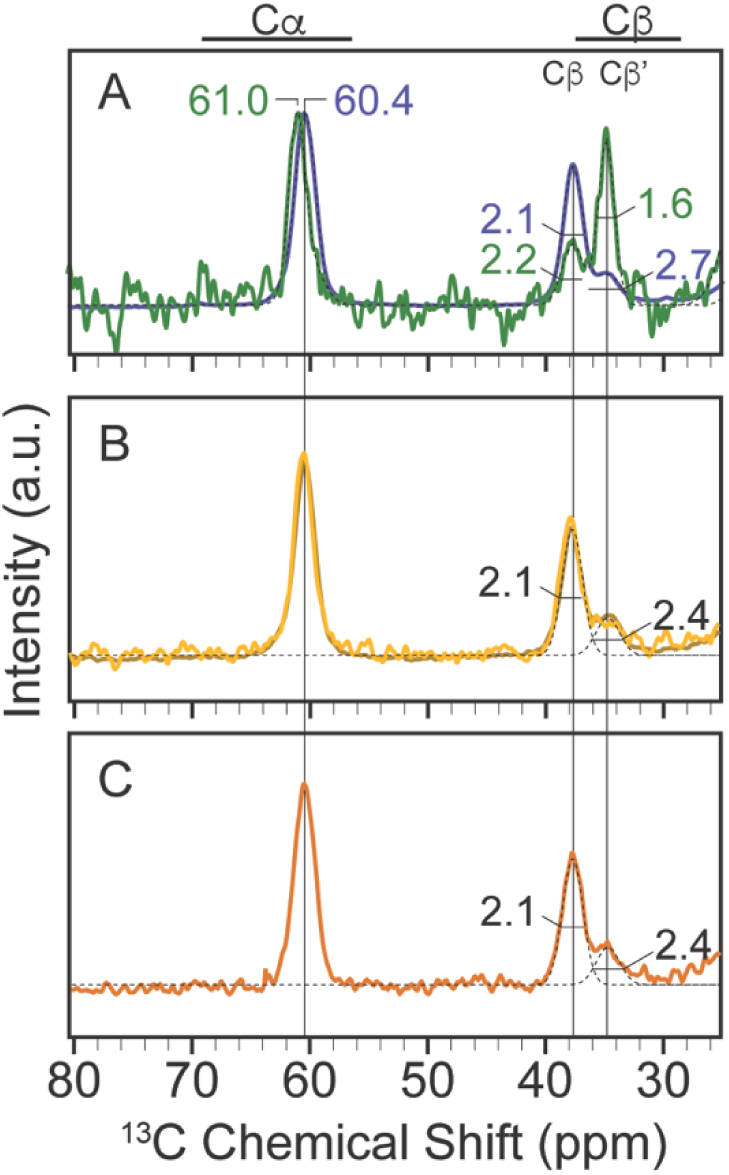
Intact cells selectively amplify and increase the molecular order of Ac-α-syn(A53T) polymorph B. A) NCOCx spectra of co-added in cell propagated ^13^C valine, ^15^N histidine labeled Ac-α-syn(A53T) fibrils (green) overlayed with the spectra of the de novo Ac-α-syn(A53T) fibrils used as seeds for cellular propagation (blue). Peak centers are annotated for the V49Cα resonance and peak widths (FWHM) are annotated for V49Cβ and V49Cβ’peaks. B) NCOCx spectrum of pre-formed Ac-α-syn(A53T) fibrils diluted into cellular lysates. C) NCOCx spectrum of Ac-α-syn(A53T) fibrils seeded by de novo Ac-α-syn(A53T) fibrils and propagated in cellular lysates. Individual Gaussians for V49Cβ and V49Cβ’ are shown as black dotted lines. NMR spectra were recorded at a field strength of 14 T (600 MHz) and MAS frequencies of 12 kHz at 104 K.

The apparent peak center for the Cα resonance was shifted 0.6 ppm in the in cell propagated fibrils relative to the *de novo* fibril seeds (**Figure 4A**). Because the V49Cβ and V49Cβ’ peak centers were unchanged, this difference suggested that the V49Cα and V49Cα’ resonances were not fully resolved under these experimental conditions, rather than that a new conformation had appeared. Consistent with this interpretation, fitting the Cα peak to a sum of two Gaussian peaks with peak centers fixed at 60.3 ppm and 61.1 ppm recapitulated both the observed change in the Cα peak position and the relative populations obtained from the V49Cβ and V49Cβ’ peaks (**Figure S8, Table S5**). These data indicate that the same two Ac-α-syn(A53T) amyloid-core conformations are present in both the seed and in-cell propagated samples, but that their relative populations invert during propagation in cells.

The inversion in population was accompanied by a change in molecular order. The linewidths of the V49Cβ peak were similar for fibrils propagated in vitro and inside cells, whereas the linewidth of the V49Cβ′ peak changed dramatically (**Figure 4A, Figure S8, Table S5**). The FWHM of V49Cβ′ narrowed from 2.7 ppm in the de novo seeds to 1.6 ppm in the in-cell propagated fibrils, a linewidth close to that of a fully rigid site under these conditions. Thus, in addition to selectively amplifying polymorph B, propagation inside cells increases its molecular order, indicating that the cell-amplified form of polymorph B is more conformationally uniform than the corresponding in vitro assembled form.

### Crowded cellular lysates fail to recapitulate the preferential amplification observed in intact cells

To determine if the increase in the population and molecular order of polymorph B in cells resulted from interactions of the amyloid fibrils with cellular constituents, we diluted a small amount of pre-formed isotopically labeled Ac-α-syn(A53T) fibrils into concentrated whole-cell lysates. In these samples, we matched the ratio of the Ac-α-syn-A53T (on a monomer-to-monomer basis) to cellular material with that of the electroporated biosensor cell samples, preserving the stoichiometry of the cellular constituents to fibrils. Cellular lysates did not alter the chemical shifts, relative polymorph populations, or the peak widths of the 1D NCOCx spectra of V49 (**Figure 4B, Table S4**). Therefore, the increase in the population of polymorph B was not the result of interactions with or remodeling by cellular constituents and the higher molecular order of in cell propagated polymorph B was not the result of either binding of cellular constituents or macromolecular crowding.

To determine whether propagation in the presence of cellular constituents and macromolecular crowding could reproduce the cellular outcome, we added a small amount of pre-formed Ac-α-syn(A53T) fibrils (1% monomer/monomer) to 70 µM monomeric isotopically labeled Ac-α-syn(A53T) in the presence of concentrated whole-cell lysates, again matching the stoichiometry of the in cell propagated samples. The resulting lysate-propagated fibrils also showed no change in chemical shifts, relative polymorph populations, or peak widths at V49 (**Figure 4C, Table S4**). Therefore, propagation in crowded cellular lysates faithfully preserves the seed ensemble rather than reproducing the selective amplification and increased molecular order observed in intact cells. Together, these two experiments show that neither cellular constituents nor macromolecular crowding are sufficient to account for the structural changes observed during propagation in cells. Instead, selective amplification and increased ordering of polymorph B are emergent properties of propagation within the organized intracellular environment of intact, viable cells.

### Flanking regions that are intrinsically disordered in purified fibrils are remodeled inside cells

To determine whether the conformational preferences of the disordered regions flanking the amyloid core are influenced by the cellular environment, we collected 2D ^13^C–^13^C DARR correlation spectrum of ^13^C valine labeled fibrils that were propagated inside cells, as well as pre-formed fibrils that were either diluted into concentrated cellular lysates or propagated in the presence of concentrated cellular lysates with 1% *de novo* seeds. In *de novo* fibrils formed *in vitro*, substantial cross-peak intensity was observed in the α-helical region of the spectrum, indicating this disordered region preferentially sampled α-helical backbone dihedral angles (**Figure 1A, 1D**). In striking contrast, fibrils propagated inside cells showed no detectable cross-peak intensity in the α-helical region of the 2D ^13^C-^13^C DARR spectrum (**Figure 5A, 5E**, light and dark green). By comparison, the relative cross peak intensity in the α-helical region of fibrils diluted into cellular lysates **(Figure 5B, 5E**, yellow) or propagated in the presence of cellular lysates **(Figure 5C, 5E**, orange) was similar to that of purified *de novo* fibrils (**Figure 1A, 5E**, red). Thus, propagation inside intact cells substantially alters the conformational preferences of the amino-terminal flanking region of Ac-α-syn(A53T) fibrils, whereas propagation in crowded lysates faithfully preserves the in vitro ensemble. Like the amyloid core, neither cellular constituents nor macromolecular crowding are sufficient to reproduce the structural changes observed in intact cells. Instead, remodeling of the flanking disordered region requires the organized intracellular environment of viable cells.

**Figure 5:**
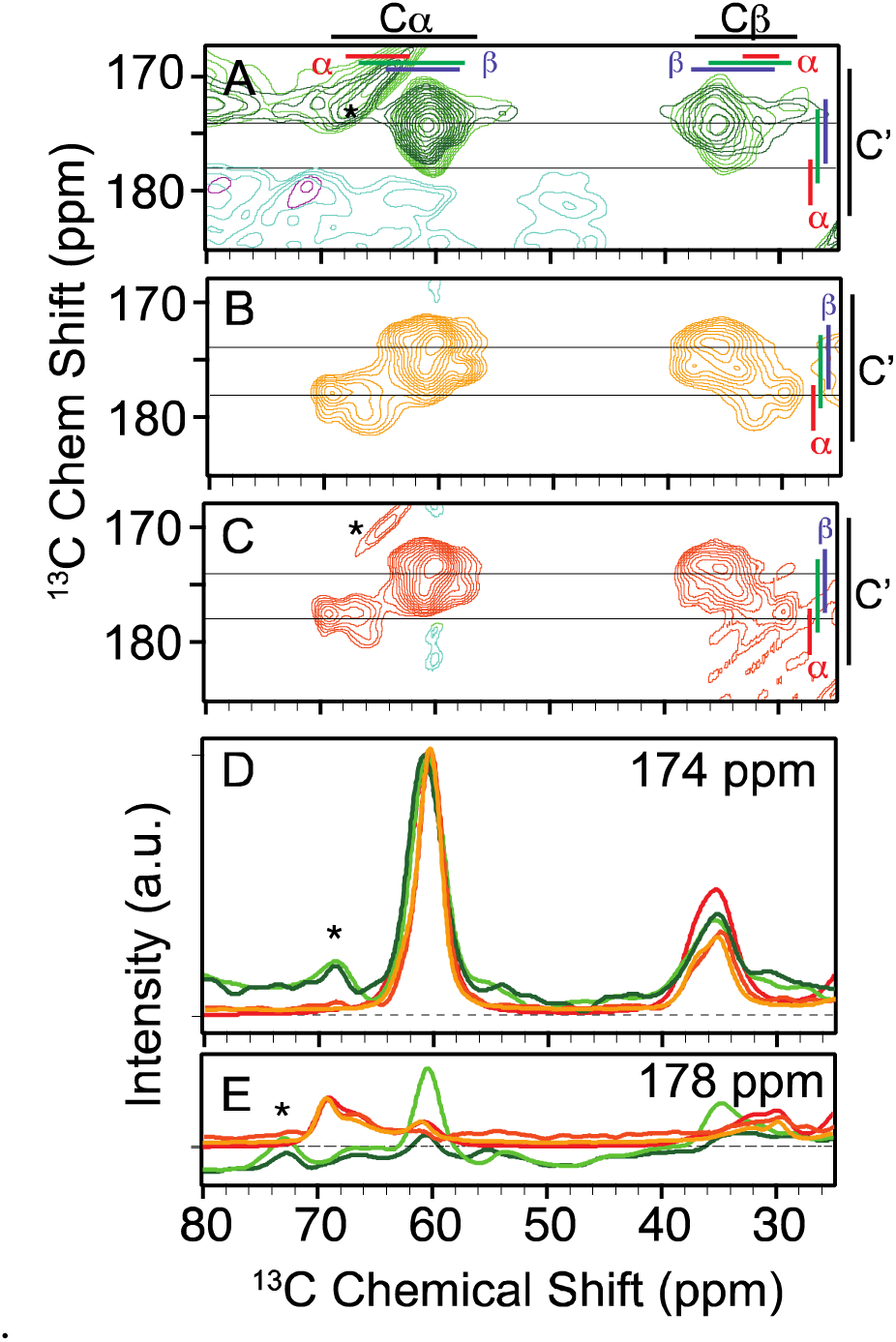
**A**) 2D ^13^C–^13^C DARR spectra (10 ms mixing) of two independent samples (light green, dark green) of ^13^C valine labeled Ac-α-syn(A53T) assembled into the amyloid conformation inside intact viable HEK293 cells. Average +/- two standard deviation of the database chemical shift values are marked for each atom (black) as well as for the α-helical (red), random coil (green), and β strand (blue) secondary structural elements. **B**) 2D ^13^C–^13^C DARR spectra of pre-formed ^13^C valine labeled Ac-α-syn(A53T) fibrils diluted in HEK293 cells lysates. **C**) 2D ^13^C–^13^C DARR spectra of ^13^C valine labeled Ac-α-syn(A53T) assembled into the amyloid conformation in HEK293 cells lysates by 1% seeding with sonicated pre-formed fibrils. One-dimensional slices from the 2D spectra of in cell amplified fibrils (dark green, light green), de novo formed amyloid seeds (shown in Figure 1A) (red), pre-formed fibrils diluted into cellular lysates (yellow), and seeded in lysate amplified fibrils (orange) at **D**) 174 ppm and **E**) 178 ppm. Spinning sidebands are marked with a star (*). NMR spectra were recorded at a field strength of 14 T (600 MHz) and a MAS frequency of 12 kHz at 104 K.

## Discussion

Here, we show that intact cells do not simply propagate the amyloid conformational ensemble present in vitro, but actively reshape it. Using DNP-assisted MAS NMR and cryo-EM, we found that the seed ensemble of Ac-α-syn(A53T) fibrils is faithfully preserved during propagation in vitro and in concentrated cellular lysates. In contrast, propagation inside intact cells produces a distinct outcome: the minority amyloid-core polymorph in vitro becomes the dominant polymorph in cells, this cell-amplified form has increased molecular order, and the amino-terminal flanking region has different conformational preferences. Thus, the biologically relevant fibril ensemble cannot be inferred from purified or lysate systems alone.

These data indicate that two distinct mechanisms operate during propagation in cells. In the amyloid core, the same two conformations are present in vitro and in cells, but their relative populations invert, consistent with selective amplification from a pre-existing ensemble (*57, 58*). In the amino-terminal flanking region, the conformational preferences themselves change, consistent with remodeling during propagation (*59*). Thus, intact cells both select among pre-existing conformers and remodel flexible regions. Notably, neither effect is reproduced in concentrated lysates, indicating that neither macromolecular crowding nor the simple presence of cellular constituents is sufficient to account for the structural changes observed in cells. Instead, both selective amplification and remodeling require the organized intracellular environment of viable cells.

Our results also clarify the distinct information provided by NMR and cryo-EM. Chemical shifts and dihedral-angle predictions link the NMR peaks to the two cryo-EM polymorphs, whereas linewidths and the fraction of particles usable for 3D reconstruction report on molecular order. Once this correspondence is established, NMR peak intensities directly report the relative populations of the two polymorphs in the frozen sample under cryogenic conditions comparable to those used for cryo-EM. Strikingly, these populations are not reflected in the cryo-EM 2D class particle counts, indicating that the number of particles assigned to a polymorph on the grid does not directly report its abundance in the sample. More broadly, these data show how quantitative spectroscopy and cryo-EM provide complementary views of the same ensemble: cryo-EM resolves atomistic structures of ordered states, whereas NMR provides access to ensemble populations and to conformational preferences in flexible regions unresolved by diffraction-based approaches.

These findings have direct implications for how amyloid structures are interpreted in biological settings. Structure-based ligand discovery and mechanistic studies typically rely on in vitro amplified fibrils, yet the dominant species in cells can differ in both population and molecular order. In our system, the polymorph that is minor and less ordered in vitro becomes selectively amplified and more ordered in cells, while the flanking region is remodeled into a distinct conformational ensemble. Together, these observations suggest that structural models derived from purified fibrils may not capture the most prevalent conformations in relevant biological contexts. More broadly, DNP-enhanced in-cell NMR fills an important gap in structural biology by enabling direct characterization of conformational ensembles in situ, including both ordered and intrinsically disordered regions.

## Acknowledgements

D.L. was supported by NIH MB T32 GM008297. S.K. was supported by UT Dallas and the Cecil and Ida Green Foundation via The Green Fellow’s Program. This work was supported by grants from the National Institute of Health [NS111236, NS134921, and NS143938] to K.K.F. We thank the Structural Biology Laboratory at UT Southwestern Medical Center which is partially supported by grant RP220582 from the Cancer Prevention & Research Institute of Texas (CPRIT) for cryo-EM studies. CEMF is supported by a grant from the Cancer Prevention & Research Institute of Texas (RP220582).

## Data and materials availability

The cryo-EM maps have been deposited in the Electron Microscopy Data Bank (EMDB) under accession codes EMD-45650 and EMD-45651. The corresponding atomic coordinates have been deposited in the Protein Data Bank (PDB) under accession codes 9CKK and 9CKL.

## Material and Methods

### Recombinant protein expression and purification

#### Protein expression

The genes for the A53T and A53T-N122C mutant form of α-syn in pET28b vector were generated by site-directed mutagenesis using Q5 high fidelity DNA polymerase (New England Biolabs) and mutation was confirmed by DNA sequencing. Amino-terminally acetylated (Ac) α-Syn(A53T) was produced by co-expression of α-syn(A53T) and the yeast N-acetyltransferase complex B (NatB) in *E. Coli* BL21-Gold DE3(Agilent) as described (*60*) ( Bell, Rosie, et al.,2022). Co-transfected cells were grown under antibiotic selection (50 µg/mL kanamycin and 34 µg/mL chloramphenicol) with aeration at 37 °C. Natural abundance proteins were expressed on 2xYT media (16 g/L tryptone, 10 g/L yeast extract 5 g/L NaCl). Valine and histidine-labeled proteins were expressed in natural abundance M9 media (48 mM Na_2_HPO4, 22 mM KH_2_PO4, 9 mM NaCl, adjusted to pH 7.4 by the addition of NaOH, 4 g/L glucose, 1 g/L NH_4_Cl, 10 mg/L FeSO4, 2 mM MgSO_4_, 100 mM CaCl_2_, 10 mg/L thiamine) supplemented with 100 mg/L of each amino acid except valine and histidine.

Expression of Ac-α-syn(A53T) and Ac-α-syn(A53T-N122C) proteins was induced by the addition of 1 mM Isopropyl β-D-1-thiogalactopyranoside (IPTG) to cells at mid-log phase (OD_600_ of 0.8-1.0). Cells were cultured for 4 hours at 37 °C. For site-specific isotope labeling ^13^C_5_ valine and ^15^N_3_ histidine (Cambridge Isotope Laboratories, Tewksbury, MA) isotope-enriched Ac-α-syn(A53T) was expressed in M9 minimal media and added before IPTG induction. Briefly, a preculture of *E.coli* bacterial BL21-Gold DE3 cells with α-syn(A53T) and NatB plasmid was grown overnight at 37°C in 300 mL M9 media supplemented with 4 gm/L glucose, 1 gm/L NH_4_Cl, 50 µg/mL Kanamycin and 34 µg/mL chloramphenicol. At the absorbance OD_600_ of 2.2-2.8, cells were collected by centrifugation and washed with 1x M9 salt and resuspended in 1 L fresh M9 minimal media containing natural abundance 4 gm/L glucose, 1 gm/L NH_4_Cl and 100 mg/L of all amino acids (Sigma) except valine and histidine. 100 mg/mL of ^13^C_5_ valine and ^15^N_3_ histidine labeled amino acid were added to the above mixture and cells were grown at 37 °C for 90 minutes, followed by induction by addition of 1 mM IPTG at the OD_600_ of 0.8-1.0. After 4 hours, cells were collected by centrifugation at 4000 x g for 20 minutes and stored at -80 °C.

#### Purification

Purification was performed as described previously (Kloepper, K. D. et al. 2006. , (*51*). Briefly, pelleted cells were resuspended in 160 mL of lysis buffer (20 mM Tris, 1 mM EDTA and 0.1% Triton X, pH 8.0) and incubated at 37 °C in the water bath. After 30 minutes, 800 μL of 2 M MgCl_2_ and 1.6 mL 1 M CaCl_2_ were added to the lysed cells along with 2.5 μL DNase (New England Biolabs) and OmniCleave endonuclease (Biosearch Technology) for nucleic acid digestion. This mixture was incubated at 37 °C for 1 hour with shaking at 200 rpm. Afterward, 1.6 mL of 0.5 M EDTA was added to the mixture to remove excess metal. Cell suspension was centrifugated at 4200 x g for 15 minutes, and 6 parts of the supernatant were mixed with 1 part of 5 M NaCl and boiled for 15 minutes in the water bath. Following another 20-minute centrifugation at 30000 x g, an equal volume of saturated ammonium sulphate (50% *v/v*) was added to the supernatant and the solution was stirred overnight at 4 °C. The precipitated Ac-α-syn(A53T) was resuspended in buffer A (20 mM NaCl, 20 mM Tris, pH 8.0). The whole mixture was dialyzed for 16-20 hours against 4 L buffer A. The dialyzed solution was filtered with 0.22 μM filter (Corning) and loaded on to a Q-Sepharose column (GE healthcare), that was pre-equilibrated with buffer A. Ac-α-syn(A53T) protein fractions were eluted with a gradient (0-80%) of buffer B (20 mM Tris, 2 M NaCl, pH 8.0). Fractions containing α-syn were concentrated to ∼1 mM with an Amicon centrifugal filter (Millipore, Sigma) and injected to superdex 75 hi-scale 26/40 size exclusion column (GE healthcare). Pure monomeric Ac-α-syn(A53T) protein fractions were pooled and purity was determined by 10% sodium dodecyl sulphate polyacrylamide gel electrophoresis (SDS-PAGE). All chromatography was performed on the Biorad NGC chromatography system (Biorad). Protein concentration was determined by UV absorption at 280 nm using a theoretical extinction coefficient of 5960 M^-1^ cm^-1^. Acetylation and protein molecular weight were confirmed by LC/MS with a Q-TOF mass spectrometer.

#### Segmentally isotopically labeled protein expression and purification

To express isotopically-labeled recombinant *N*^α^-acetylated α-syn(1–29)-Cfa_N_ construct, an overnight pre-culture of BL21(DE3) cells carrying simultaneous plasmids encoding α-syn(1–29)-Cfa_N_ and yeast NatB was used to inoculate 4 liters of M9 media with ^13^C_5_-valine was added at a final concentration of 100 mg/L containing 50 mg/L kanamycin and 34 mg/L chloramphenicol.. Cells were grown at 37 °C with shaking until their *A*_600_ reached 0.6–0.8; the protein expression was then induced by adding isopropyl 1-thio-β-D-galactopyranoside (IPTG) to a final concentration of 1 mM. Cells were harvested 4 hours after induction by centrifugation at 4000 × *g* for 15 min at 4 °C.

Natural-abundance recombinant Cfa_C_-α-syn(C30-140, A53T) was expressed in LB media (10 g/L tryptone, 5 g/L yeast extract, 10 g/L NaCl). Cells were grown at 37 °C with shaking until their *A*_600_ reached 0.6–0.8, and protein expression was induced by the addition of IPTG to a final concentration of 1 mM. Cells were collected 4 h later by centrifugation at 4,000 × *g* for 15 min at 4 °C.

Proteins from constructs 1 and 2 were purified using nickel resin. Cell pellets were lysed by resuspension in 20 mL of binding buffer (100 mM NaH_2_PO_4_, 500 mM NaCl, 4 mM TCEP, pH 8.0) per liter of growth. Protease inhibitor tablets were added. The cell suspension was sonicated 6 times (pulse: 60% amplitude, 30 s, 0.5 s on, 0.5 s off). Lysates were centrifuged for 20 min at 30,000 × *g* to remove insoluble cellular debris. Cleared lysates were filtered using 0.2 µM filter then passed through a 15-mL Nickel column using Biorad NGC system. The column was washed with 10-15 column volumes (CV) of binding buffer (100 mM NaH_2_PO_4_, 500 mM NaCl, 2 mM TCEP, pH 8.0). It was then washed with 10-15 CV of wash buffer (100 mM NaH_2_PO_4_, 500 mM NaCl, 10 mM imidazole, 2 mM TCEP, pH 8). The protein was eluted in elution buffer (100 mM NaH_2_PO_4_, 500 mM NaCl, 450 mM imidazole, 2 mM TCEP, pH 8). Purity was assessed by SDS/PAGE. Protein concentration was quantitated using theoretical extinction coefficients of 26930 M^−1^⋅cm^−1^ for construct 1 and 7450 M^−1^⋅cm^−1^ for construct 2.

#### Split-Intein Ligation and Purification

The purified split-intein constructs were ligated at room temp overnight with a molar excess of 1:1.5 (construct 1:construct 2) in ligation buffer [50 mM Tris, 300 mM NaCl, 2 mM Tris(2-carboxyethyl)phosphine (TCEP), 0.5 mM EDTA, pH 7.4]. The final concentrations of constructs 1 and 2 were 25 µM and 30 µM, respectively. The reaction mixture was left at 4 °C overnight. It was then concentrated 5-fold using a 10-kDa molecular mass cutoff Amicon spin concentrator. The reaction mixture was then incubated with Ni-NTA resin for an hour at room temperature. The resin was transferred to a PD-10 column and washed with 1 CV of binding buffer. The flowthrough and the wash were collected since they contain the Ac-α-syn(A53T) that is untagged. The protein solution was concentrated 10-15 fold before proceeding to the desulfurization step.

Desulfurization was done following a previously published protocol, which contains important safety precautions (*61*). Briefly, pH-adjusted TCEP was added to the protein solution to bring up the concentration of TCEP to 100 mM. Then the radical initiator solution (100 mM VA-044 in 6 M guanidine hydrochloride, 50 mM sodium phosphate, pH 7.2) was added to a final concentration of 6 mM. The following steps are all done in a fume hood. Methylpropane-2-thiol (or t-butyl mercaptan) was added in the hood to a final concentration of 400 mM. The reaction was left on an orbital shaker for 2 hours at 600-800 rpm at 37 °C. The protein was then isolated in the hood using PD10 columns equilibrated with the buffer of interest.

At this step we had Ac-α-syn(A53T) and an 11-kDa byproduct of the ligation reaction (the last 111 amino acids of synuclein). Therefore, after the desulfurization reaction, the protein was concentrated and loaded onto a Superdex 75 size exclusion column. The fractions containing isolated full-length Ac-α-syn(A53T) were pooled together.

#### Maleimide Labeling of Ac-α-syn(A53T- N122C)

Alexa fluor 647 maleimide (AF^647^) (Invitrogen) was site-specifically attached to Ac-α-syn(A53T- N122C) protein at position 122. Before labeling with AF^647^, the protein was incubated in degassed labeling buffer (100 mM NaHPO_4_, 5 mM KCl, 15 mM HEPES, pH 7.0) with a 10-fold excess of TCEP (Gold Biotechnology). The excess of TCEP was removed by filtration over a NAP-10 column (Sephadex G-25, Cytiva). The maleimide dye was added to the protein at a 4:1 molar ratio and the reaction mixture was incubated in the dark at 25 °C for 4 h. The labeling reaction was quenched by addition of excess of β-mercaptoethanol and free fluorophore was removed by using a PD-10 desalting column (Cytiva). The labeling efficiency was calculated as described (*62*) and samples were concentrated with Amicon centrifugal filters.

### Preparation of *in vitro* fibrils

Ac-α-syn(A53T) *de novo* fibrils were prepared as described (*41*). Briefly, monomeric protein was diluted to 300 μM in Dulbecco’s phosphate-buffered saline (DPBS, Gibco) and incubated at 37 °C for 5 days with constant agitation at 1000 rpm in an Eppendorf orbital thermomixer. First generation fibrils were prepared by adding 1% (*mol/mol*) of *de novo* fibrils to the monomer at the start of the polymerization reaction. For DNP NMR studies, fibrils were collected by centrifugation at 200,000 x g for 1 hour and the supernatant was removed. The pellet was resuspended in perdeuterated 1x PBS buffer containing a final composition of either 60% *d_8_*-glycerol, 30% D_2_O, 10% H_2_O pH 7.4, and 7 mM AMUPol or 15% *d_8_*-glycerol, 75% D_2_O,10% H_2_O pH 7.4 and 7 mM AMUPol. The sample was packed into a 3.2 mm sapphire rotor (Bruker) and stored at -80 °C.

### Transmission electron microscopy

Ac-α-syn(A53T) fibril samples (5 μL) were loaded onto a glow-discharged carbon-coated electron microscopy grid. After 1 min, the grid was washed 3 times with 5 μL distilled water and 5 μL uranyl acetate (2% in aqueous solution) was applied to the grid for 1 min. Filter paper was used to remove excessive stains. The samples were imaged using an FEI Tecnai G2 Spirit Biotwin electron microscope.

### Cryo-EM sample preparation and data acquisition

The Ac-α-syn(A53T) fibril sample was the applied to Quantifoil 300-mesh R1.2/1.3 grids (Quantifoil, Micro Tools GmbH, Germany) that were pre-treated using a Pelco EasiGlow instrument (Ted Pella). The grid was flash-frozen into liquid ethane using a Vitrobot Mark IV (Thermo Fisher Scientific), with the following settings: blot time 3.5 s, relative humidity 95%, and 4 °C.

Grid screening and data collection were performed at the UTSW Cryo-Electron Microscopy Facility (CEMF). After screening, the best grid was used for a 24-hour data collection on a Titan Krios microscope (Thermo Fisher Scientific) equipped with the post-column BioQuantum energy filter (Gatan) and a K3 direct electron detector (Gatan). Cryo-EM data were collected using SerialEM (*63*) in a super-resolution mode with a 20 eV energy filter slit in CDS mode. 7020 movies were acquired in super-resolution mode, at a pixel size of 0.42 Å. The accumulated total dose was 50 e^-^/Å^2^ for each movie stack and it was fractionated into 50 frames. The defocus range of the images was set to be −0.9 to −2.2 μm.

#### Cryo-EM data processing

Cryo-EM data were processed using Relion (*64–66*). Motion correction was performed using MotionCor2 (*67*) and the movies were averaged into single images with a binning factor of 2, resulting in a pixel size of 0.84 Å/pixel. The CTF parameters were calculated using Gctf (*68*). Fibril auto-picking was done using Topaz (*69*) within the Relion pipeline, and 2,266,580 particles were extracted at 800 box size with an inter-box distance of 38 Å, down sampled to 2.52 Å/pixel for 2D classification. Two types of fibrils were observed in the 2D classes based on the distinct features. The crossover distance for each fibril type was estimated from the 2D classes and was used in relion_helix_inimodel2d to generate the initial model. For Polymorph A, 813,225 particles were selected and re-extracted at 300 box size, followed by 3D classification, 3D refinement, and CTF refinement. 85,236 particles were used to reconstruct the final density map, and the reported resolution was 2.21 Å by Relion post-processing. The polymorph A fibril structure adopts a C2_1_ screw symmetry. The refined helical twist is 179.61 degrees, and the helical rise is 2.40 Å. For polymorph B, 983,759 particles were selected and re-extracted at 300 box size. After multiple rounds of 3D classification, 11,326 particles were selected for 3D refinement to generate the final density map. The resolution of this map was 2.68 Å as estimated by Relion post-processing. Polymorph B structure adopts a C2 symmetry. The refined helical twist is -1.21 degrees, and the helical rise is 4.81 Å.

#### Model building and refinement

For both Polymorph A and Polymorph B structures, an initial model was first built using Model Angelo (*70*) in the Relion pipeline, followed by manual building in Coot (*71*) and Chimera (*72*). The models were then refined with Phenix Real-space refinement (*73*). The statistics of the model validations are summarized in Table S2.

### Cellular NMR sample preparation

To introduce isotopically labeled α-syn(A53T) into mammalian cells, we used a modified protocol based on previous work (*43*). HEK293T α-syn(A53T)-CFP/YFP biosensor cell lines were grown in DMEM (Gibco) with 10% FBS (Gibco) and 0.5% GlutaMAX (Gibco) at 37 °C in 5% CO_2_ humidified incubator on a 10 cm dish to confluency. Cells were washed with 1x PBS (Gibco) and harvested using Tryp-LE express (Gibco). 48 × 10^6^ cells were collected by centrifugation (233 x g, 5 min) and resuspended in 1 mL electroporation buffer (EPB) 100 mM NaHPO_4_, 5 mM KCl, 15 mM HEPES, 15 mM MgCl_2_, 2 mM reduced glutathione and 2 mM ATP, pH 7.0. Cells suspension in EPB were collected by centrifugation at 233 x g for 5 min and supernatant was discarded. Cell pellets (∼48 x 10^6^ cells) were resuspended in 1.1 mL EPB containing a final concentration of 800 μM monomeric α-syn(A53T). α-syn(A53T) was introduced into cells by electroporation. 100 μL aliquots of the cells and α-syn(A53T) mixture in EPB were transferred to electroporation cuvettes (Lonza) and electroporation was done on the Lonza 4D nucleofector system by using the CM-130 pulse program. Cells were pulsed two times and gently mixed between the pulses. After electroporation, 500 μL of cell culture media was added to each cuvette and mixed with the cells using a Pasteur pipette and all electroporated cells were added to 10 mL of prewarmed cell culture medium in a 100 mm diameter dish (Corning) and incubated for 5 hours at 37°C with 5% CO_2_.

#### Seeding

After electroporated cells recovered for 5 hours, adherent cells were rinsed with 1 x PBS and detached using Tryp-LE express. For in cell seeding, fibrils were sonicated for 15 minutes at 4 °C in a water bath using an ultrasonicator (QSonica, Q700) with pulses of 3 sec on/2 sec off and 30% amplitude. Sonicated Ac-α-syn(A53T) fibrils (11.5 µL, 100 _µ_M) were diluted in 738 μl of Opti-MEM. In the second tube, 30 μl of Lipofectamine (Invitrogen, 11668019) was added with 720 μl of Opti-MEM and incubated at RT for 5 minutes. The 1.5 mL mixture of fibrils and lipofectamine was incubated at RT for 30 minutes before dropwise addition to the cell suspension (4-4.5 x 10^6^ cells) in 10 mL DMEM media with 10% FBS and 1% Penicillin-Streptomycin. The final concentration of the sonicated Ac-α-syn(A53T) fibrils was 100 nM. Cells were incubated at 37 °C with 5% CO_2_ and harvested after 24 hours.

### Microscopy

For microscopy, samples were treated identically except that Ac-α-syn(A53T-N122C)-AF^647^ was introduced into cells at the electroporation step instead of Ac-α-syn(A53T). After the 24-hour incubation step, cells were washed with 1x dPBS (Thermofisher, 14040133) and stained with 2.5 μg/mL cell mask orange plasma membrane stain (Invitrogen, C10045) for 5 min at 37 °C. Cells were quickly washed 3 times with cold 1x dPBS and fixed in 4 % (*w/v*) paraformaldehyde (PFA) for 15 minutes at room temperature. After washing once in 1x dPBS cells were stored in 1x live cell imaging solution (Invitrogen, A59688DJ) for microscopy. Confocal images were taken at 63x magnification on a Zeiss LSM 880 Confocal Microscope by using the excitation wavelengths of 405, 561, and 633 nm for CFP, cell mask orange and AF^647^ respectively. Images were analyzed using Fiji software.

### FRET experiments

For the FRET seeding assay, cells were harvested after 24 hours and fixed with 2% paraformaldehyde (Electron Microscopy Sciences) for 10 minutes. After centrifugation cell pellet was resuspended in 2 mL cold 1x PBS for flow cytometry analysis. Three additional control cell lines (HEK293T cells, α-syn(A53T)-YFP and α-syn(A53T)-CFP transduced cells) were used in this assay. The sample was analyzed by LSRII Fortessa flow cytometer (BD Biosciences), where the FRET signal was detected using the CFP-specific 405 nm laser with 525/50 bandpass filter. CFP and YFP are excited by 405 nm and 488 nm lasers with 450/50 and 525/50 nm bandpass filters, respectively. For sorting, cells were seeded with 100 nM Ac-α-syn(A53T) fibril for 24 h and harvested. Cells pellet mixed with 1 mL cold Flow buffer (1X Hank Balanced salt solution, Ca^++^/Mg^++^ free, 1% FBS and 1 mM EDTA) and filtered through cellTrics filter (Sysmex). FRET negative and positive cells were sorted at 4 °C using FACSArias by using the CFP-specific 405 nm laser with a 525/50 bandpass filter.

#### Growth Assay

600,000 cells were plated in 10 cm dishes containing 10 mL warm media (DMEM). Media was changed after cells had adhered overnight and cells were incubated for 4-5 days. Confluency was monitored daily at 3 independent areas of each dish with an inverted light microscope (Olympus CKX53 Tokyo, Japan). Confluency was determined using ImageJ.

#### Annexin-PI assay

The Annexin V-Alexa Fluor 647 conjugate and propidium iodide (PI, Invitrogen) combination was employed to quantitate apoptosis following the manufacturer’s instructions. Briefly, cells were seeded with 100 nM Ac-α-syn(A53T) *de novo* fibril and incubated for 24-72 h. Post-treatment, cells in 10 mL culture media were stained with 2.5 μL annexin V-Alexa Fluor 647 and 1 μL PI for 15 min in the dark before cytometry analysis with a LSRII Fortessa flow cytometer (BD Biosciences). The FRET signal was detected using CFP specific 405 nm laser with a 525/50 bandpass filter. We plotted the FRET versus CFP bivariate plot and introduced a gate to separate the FRET-positive and negative cell populations. To detect apoptosis in the FRET positive population, Annexin V-Alexa Fluor 647 and PI fluorescence emission was measured using the 670/20 nm bandpass and 660/20 long-pass filter with excitation wavelengths of 633 and 532 nm, respectively. A total of 30,000 events per sample were recorded, and compensation was set up using unstained cells and cells stained with a single-color dye only. Data were analyzed using FlowJo 10.9.0 software.

### NMR sample preparation of in cell propagated fibril

For the intact cell sample, pelleted cells were washed with 1 mL of perdeuterated 1x PBS (88% D_2_O, 10% H_2_O, pH 7.4). After centrifugation (233 x g, 5 min), the 50 μL cell pellet was mixed with 50 μL of perdeuterated 1x PBS containing AMUPol (Cortecnet, USA) and 18 μL *d_8_*-^13^C-depleted glycerol (Cambridge Isotope Labs). The 118 μL cell suspension had a final sample matrix of 15:75:10 *d_8_*-^13^C-depleted glycerol:D_2_O:H_2_O (*v*/*v*/*v*) with 5 mM AMUPol. Cells were packed in a 3.2 mm sapphire rotor (Bruker) by centrifugation at 100 x g for 1 min. The rotor was sealed with a silicone plug and capped with a zirconia drive cap. The rotor was frozen for 6 hours at a controlled rate of 1 °C/min in “Cool Cell LX” (Corning) in the −80 °C freezer before transfer to liquid nitrogen until NMR analysis.

The fibril-containing fraction of one sample of in cell propagated fibrils was increased to improve sample signal to noise. To do so, pelleted cells were resuspended in 500 μL of perdeuterated 1x PBS (88% D_2_O, 10% H_2_O pH 7.4). Cells were lysed in 500 μL buffer by three freeze-thaw cycles in liquid nitrogen. The insoluble high molecular weight cellular material was collected by centrifugation at 200,000 x g for 10 minutes. The 25 μL pellet was mixed with 25 μL of perdeuterated 1x PBS containing AMUPol (Cortecnet, USA) and 9 μL *d_8_*-^13^C-depleted glycerol. The insoluble portion was collected in a 3.2 mm rotor and the final sample composition was 15:75:10 *d_8_*-glycerol:D_2_O:H_2_O (*v*/*v*/*v*) with 5 mM AMUPol.

For the dilution of pre-formed VH-labeled Ac-α-syn(A53T) fibrils into cellular lysates, 50 μL cell pellet lysate having fibril was mixed with 50 μL of perdeuterated 1x PBS (85% D_2_O + 10% H_2_O, pH 7.4) containing AMUPol (Cortecnet, USA) and 18 μL of *d_8_*-^13^C-depleted-glycerol. The 118 μL cell suspension had a final composition of 23 μM fibril, 5 mM AMUPol with 15% (*v/v*) *d_8_*-^13^C-depleted-glycerol, 75% (*v/v*) D_2_O, and 10% (*v/v*) H_2_O.

For the seeding amplification of VH-labeled Ac-α-syn(A53T) monomer in cellular lysates of sensor cells that had never been exposed to amyloid fibrils, a 50 μL cell pellet was lysed by three freeze thaw cycles in the presence of 1x protease inhibitor (Sigma-Aldrich, 11836170001) and VH-labeled Ac-α-syn(A53T) monomer was diluted into the lysate to a concentration of 75 μM. Sonicated, pre-formed VH-labeled Ac-α-syn(A53T) fibrils were added at a calculated monomer concentration of 1% (0.75 µM) relative to the exogenously added Ac-α-syn(A53T) (∼0.3% relative to total α-syn content of the sample). The mixture was incubated for 24 hours at 37 °C in a 5% CO_2_ atmosphere. After incubation, we added 500 μL of perdeuterated 1x PBS (85% D2O + 10% H2O, pH 7.4) to the sample and collected the insoluble high molecular weight cellular material by centrifugation at 200,000 x g for 10 minutes. The ∼25 μL pellet was mixed with 25 μL of perdeuterated 1x PBS containing AMUPol (Cortecnet, USA) and 9 μL d_8_-^13^C-depleted glycerol and transferred to a 3.2 mm rotor. The final sample composition was 15:75:10 *d_8_*-^13^C-depleted glycerol:D_2_O:H_2_O (*v/v/v*) with 5 mM AMUPol.

### NMR Spectroscopy

Rotors were transferred in liquid nitrogen directly into the NMR probe that had been previously equilibrated to 100 K (*38*). All dynamic nuclear polarization magic angle spinning nuclear magnetic resonance (DNP MAS NMR) experiments were performed on a 600 MHz Bruker Ascend DNP NMR spectrometer/7.2 T Cryogen-free gyrotron magnet (Bruker), equipped with a ^1^H, ^13^C, ^15^N triple-resonance, 3.2 mm low temperature (LT) DNP MAS NMR Bruker probe (600 MHz). The sample temperature was 104 K and MAS frequency was 12 kHz. For ^13^C cross-polarization (CP) MAS experiments, the ^13^C radio frequency (RF) amplitude was fixed at 60 kHz and an ^1^H RF amplitude was 72 kHz. The 90° ^1^H pulse was 100 kHz, the 90° ^13^C pulse was 62.5 kHz, and ^1^H TPPM at 85 kHz for decoupling with phase alternation of ± 15° during acquisition of ^13^C signal. ^13^C-^13^C 2D correlations were measured using 10 ms DARR mixing with the ^1^H amplitude at the MAS frequency. A total of 280 points in the indirect dimension were recorded with an increment of 25 µs. DARR experiments were apodized with a Lorenz-to-Gauss window function with IEN of 25 Hz and GB of 75 Hz in the *t_1_* and *t_2_*time domains. The noise level and peak height from the 2D NMR spectrum was detected by the NMRDraw software for signal-to-noise estimation. For NCOCX double CP experiment, the first CP was applied with ^1^H RF amplitude linearly swept from 51 to 101 kHz and ^15^N amplitude at 40 kHz, at a 1.8 ms contact time. The second CP was applied with an upward tangential ramp for ^13^C (tan50) RF amplitude from 24 kHz to 36 kHz, ^15^N amplitude at 42 kHz, and ^1^H constant wave decoupling at 98 kHz, at a 6 ms contact time. The spin diffusion was applied with ^1^H constant wave RF amplitude at 12 kHz for 50 ms.

#### Gaussian fitting for 1D NCOCx spectrum

For fitting of 1D NCOCx experiments, the data were processed using NMRPipe. The real part of the processed spectrum was exported using pipe2txt.tcl command. A sum of four Gaussian functions was used to fit the NCOCx spectra. The Gaussian function is defined as:

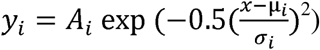

where *A_i_* is the amplitude, μ*_i_* is the mean, and σ*_i_* is the standard deviation of the i-th Gaussian function. A constant offset c was included to account for baseline correction. The total fitting function is given by:

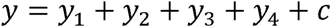

An initial guess for the parameters was inputted, and the lower and upper bounds for the parameters were set to constrain the optimization process. The built-in MATLAB function ‘lsqcurvefit’ was used to fit the Gaussian functions to the data. The peak centers for the Cα and Cα’ peaks were fixed at 61.1 and 60.3 ppm, respectively, while the peak centers for the Cβ and Cβ’ peaks were allowed to vary.

## References cited

1. M. Schweighauser et al., Structures of alpha-synuclein filaments from multiple system atrophy. Nature 585, 464–469 (2020).

2. L. Frey et al. (eLife Sciences Publications, Ltd, 2024).

3. S. Lovestam, S. H. W. Scheres, High-throughput cryo-EM structure determination of amyloids. Faraday Discuss 240, 243–260 (2022).

4. J. P. Seibyl, alpha-Synuclein PET and Parkinson Disease Therapeutic Trials: Ever the Twain Shall Meet? J Nucl Med 63, 1463–1466 (2022).

5. H. Miake, H. Mizusawa, T. Iwatsubo, M. Hasegawa, Biochemical characterization of the core structure of alpha-synuclein filaments. J Biol Chem 277, 19213–19219 (2002).

6. A. Tarutani, T. Arai, S. Murayama, S.-i. Hisanaga, M. Hasegawa, Potent prion-like behaviors of pathogenic α-synuclein and evaluation of inactivation methods. Acta Neuropathologica Communications 6, 29 (2018).

7. S. J. Wood et al., alpha-synuclein fibrillogenesis is nucleation-dependent. Implications for the pathogenesis of Parkinson’s disease. J Biol Chem 274, 19509–19512 (1999).

8. M. D. Tuttle et al., Solid-state NMR structure of a pathogenic fibril of full-length human alpha-synuclein. Nat Struct Mol Biol 23, 409–415 (2016).

9. C. Kim et al., Exposure to bacterial endotoxin generates a distinct strain of α-synuclein fibril. Sci Rep 6, 30891 (2016).

10. C. Peng et al., Cellular milieu imparts distinct pathological α-synuclein strains in α-synucleinopathies. Nature 557, 558–563 (2018).

11. F. De Giorgi et al., Novel self-replicating α-synuclein polymorphs that escape ThT monitoring can spontaneously emerge and acutely spread in neurons. Sci Adv 6, (2020).

12. S. Lovestam et al., Seeded assembly in vitro does not replicate the structures of alpha-synuclein filaments from multiple system atrophy. FEBS Open Bio 11, 999–1013 (2021).

13. D. B. Berry et al., Drug resistance confounding prion therapeutics. Proceedings of the National Academy of Sciences 110, E4160–E4169 (2013).

14. A. W. P. Fitzpatrick et al., Cryo-EM structures of tau filaments from Alzheimer’s disease. Nature 547, 185–190 (2017).

15. R. Guerrero-Ferreira et al., Cryo-EM structure of alpha-synuclein fibrils. eLife 7, e36402 (2018).

16. X. Yang, B. Wang, C. L. Hoop, J. K. Williams, J. Baum, NMR unveils an N-terminal interaction interface on acetylated-α-synuclein monomers for recruitment to fibrils. Proceedings of the National Academy of Sciences 118, e2017452118 (2021).

17. P. Kumari et al., Structural insights into α-synuclein monomer–fibril interactions. Proceedings of the National Academy of Sciences 118, e2012171118 (2021).

18. A. Peduzzo, S. Linse, A. K. Buell, The Properties of α-Synuclein Secondary Nuclei Are Dominated by the Solution Conditions Rather than the Seed Fibril Strain. ACS Chem Neurosci 11, 909–918 (2020).

19. S. F. Falsone et al., SERF protein is a direct modifier of amyloid fiber assembly. Cell Rep 2, 358–371 (2012).

20. X. Gao et al., Human Hsp70 Disaggregase Reverses Parkinson’s-Linked α-Synuclein Amyloid Fibrils. Mol Cell 59, 781–793 (2015).

21. D. Cox et al., The small heat shock protein Hsp27 binds α-synuclein fibrils, preventing elongation and cytotoxicity. J Biol Chem 293, 4486–4497 (2018).

22. E. M. Martin et al., Conformational flexibility within the nascent polypeptide-associated complex enables its interactions with structurally diverse client proteins. J Biol Chem 293, 8554–8568 (2018).

23. B. M. Burmann et al., Regulation of α-synuclein by chaperones in mammalian cells. Nature 577, 127–132 (2020).

24. S. D. Khare, P. Chinchilla, J. Baum, Multifaceted interactions mediated by intrinsically disordered regions play key roles in alpha synuclein aggregation. Current Opinion in Structural Biology 80, 102579 (2023).

25. S. M. Ulamec, D. J. Brockwell, S. E. Radford, Looking Beyond the Core: The Role of Flanking Regions in the Aggregation of Amyloidogenic Peptides and Proteins. Frontiers in Neuroscience 14, (2020).

26. R. H. Havlin, R. Tycko, Probing site-specific conformational distributions in protein folding with solid-state NMR. Proceedings of the National Academy of Sciences 102, 3284–3289 (2005).

27. K. N. Hu, R. H. Havlin, W. M. Yau, R. Tycko, Quantitative determination of site-specific conformational distributions in an unfolded protein by solid-state nuclear magnetic resonance. J Mol Biol 392, 1055–1073 (2009).

28. B. Uluca et al., DNP-enhanced MAS NMR: a tool to snapshot conformational ensembles of α-synuclein in different states. Biophysical journal 114, 1614–1623 (2018).

29. J. Kragelj et al., Spatially resolved DNP-assisted NMR illuminates the conformational ensemble of α-synuclein in intact viable cells. Proc Natl Acad Sci U S A 122, e2500367122 (2025).

30. R. Dumarieh et al., Structural Context Modulates the Conformational Ensemble of the Intrinsically Disordered Amino Terminus of α-Synuclein. J Am Chem Soc 147, 11800–11810 (2025).

31. L. Levorin et al., Isoleucine Side Chains as Reporters of Conformational Freedom in Protein Folding Studied by DNP-Enhanced NMR. J Am Chem Soc 147, 15867–15879 (2025).

32. K. K. Frederick et al., Sensitivity-enhanced NMR reveals alterations in protein structure by cellular milieus. Cell 163, 620–628 (2015).

33. W. N. Costello, Y. Xiao, K. K. Frederick, DNP-Assisted NMR Investigation of Proteins at Endogenous Levels in Cellular Milieu. Methods Enzymol 615, 373–406 (2019).

34. S. Narasimhan et al., DNP-Supported Solid-State NMR Spectroscopy of Proteins Inside Mammalian Cells. Angew Chem Int Ed Engl 58, 12969–12973 (2019).

35. D. Beriashvili et al., A high-field cellular DNP-supported solid-state NMR approach to study proteins with sub-cellular specificity. Chemical Science 14, 9892–9899 (2023).

36. R. Ghosh, Y. Xiao, J. Kragelj, K. K. Frederick, In-Cell Sensitivity-Enhanced NMR of Intact Viable Mammalian Cells. J Am Chem Soc 143, 18454–18466 (2021).

37. W. N. Costello, Y. Xiao, F. Mentink-Vigier, J. Kragelj, K. K. Frederick, DNP-assisted solid-state NMR enables detection of proteins at nanomolar concentrations in fully protonated cellular milieu. Journal of Biomolecular NMR, (2024).

38. R. Ghosh, J. Kragelj, Y. Xiao, K. K. Frederick, Cryogenic Sample Loading into a Magic Angle Spinning Nuclear Magnetic Resonance Spectrometer that Preserves Cellular Viability. J Vis Exp, (2020).

39. Y. Xiao, R. Ghosh, K. K. Frederick, In-Cell NMR of Intact Mammalian Cells Preserved with the Cryoprotectants DMSO and Glycerol Have Similar DNP Performance. Front Mol Biosci 8, 789478 (2021).

40. R. Ghosh, R. Dumarieh, Y. Xiao, K. K. Frederick, Stability of the nitroxide biradical AMUPol in intact and lysed mammalian cells. J Magn Reson 336, 107150 (2022).

41. Y. Sun et al., Cryo-EM structure of full-length α-synuclein amyloid fibril with Parkinson’s disease familial A53T mutation. Cell Res 30, 360–362 (2020).

42. J. I. Ayers et al., Different alpha-synuclein prion strains cause dementia with Lewy bodies and multiple system atrophy. Proc Natl Acad Sci U S A 119, (2022).

43. F. X. Theillet et al., Structural disorder of monomeric alpha-synuclein persists in mammalian cells. Nature 530, 45–50 (2016).

44. D. W. Sanders et al., Distinct tau prion strains propagate in cells and mice and define different tauopathies. Neuron 82, 1271–1288 (2014).

45. A. L. Woerman et al., Propagation of prions causing synucleinopathies in cultured cells. Proceedings of the National Academy of Sciences 112, E4949–E4958 (2015).

46. S. B. Prusiner et al., Evidence for α-synuclein prions causing multiple system atrophy in humans with parkinsonism. Proceedings of the National Academy of Sciences 112, E5308–E5317 (2015).

47. A. L. Woerman et al., MSA prions exhibit remarkable stability and resistance to inactivation. Acta Neuropathologica 135, 49–63 (2018).

48. J. Pauli, M. Baldus, B. van Rossum, H. de Groot, H. Oschkinat, Backbone and Side-Chain 13C and 15N Signal Assignments of the α-Spectrin SH3 Domain by Magic Angle Spinning Solid-State NMR at 17.6 Tesla. ChemBioChem 2, 272–281 (2001).

49. Y. Wang, O. Jardetzky, Probability-based protein secondary structure identification using combined NMR chemical-shift data. Protein Sci 11, 852–861 (2002).

50. M. V. Berjanskii, S. Neal, D. S. Wishart, PREDITOR: a web server for predicting protein torsion angle restraints. Nucleic Acids Res 34, W63-69 (2006).

51. J. Kragelj, R. Dumarieh, Y. Xiao, K. K. Frederick, Conformational ensembles explain NMR spectra of frozen intrinsically disordered proteins. Protein Sci, e4628 (2023).

52. K. Asanuma, K. N. Campbell, K. Kim, C. Faul, P. Mundel, Nuclear relocation of the nephrin and CD2AP-binding protein dendrin promotes apoptosis of podocytes. Proceedings of the National Academy of Sciences 104, 10134–10139 (2007).

53. F.-X. Theillet et al., Site-specific NMR mapping and time-resolved monitoring of serine and threonine phosphorylation in reconstituted kinase reactions and mammalian cell extracts. Nature protocols 8, 1416 (2013).

54. F.-X. Theillet et al., Structural disorder of monomeric α-synuclein persists in mammalian cells. Nature 530, 45–50 (2016).

55. J. M. Plitzko, B. Schuler, P. Selenko, Structural Biology outside the box-inside the cell. Curr Opin Struct Biol 46, 110–121 (2017).

56. D. Pinotsi et al., Direct Observation of Heterogeneous Amyloid Fibril Growth Kinetics via Two-Color Super-Resolution Microscopy. Nano Letters 14, 339–345 (2014).

57. J. Collinge, Prion strain mutation and selection. Science 328, 1111–1112 (2010).

58. J. Li, S. Browning, S. P. Mahal, A. M. Oelschlegel, C. Weissmann, Darwinian Evolution of Prions in Cell Culture. Science 327, 869–872 (2010).

59. N. Makarava, I. V. Baskakov, The evolution of transmissible prions: the role of deformed templating. PLoS Pathog 9, e1003759 (2013).

60. M. Johnson, A. T. Coulton, M. A. Geeves, D. P. Mulvihill, Targeted amino-terminal acetylation of recombinant proteins in E. coli. PLoS One 5, e15801 (2010).

61. B. Fauvet, H. A. Lashuel, in Protein Amyloid Aggregation: Methods and Protocols, D. Eliezer, Ed. (Springer New York, New York, NY, 2016), pp. 3-20.

62. Y. Kim et al., Efficient site-specific labeling of proteins via cysteines. Bioconjug Chem 19, 786–791 (2008).

63. D. N. Mastronarde, Automated electron microscope tomography using robust prediction of specimen movements. J Struct Biol 152, 36–51 (2005).

64. S. H. Scheres, RELION: implementation of a Bayesian approach to cryo-EM structure determination. J Struct Biol 180, 519–530 (2012).

65. R. Fernandez-Leiro, S. H. W. Scheres, A pipeline approach to single-particle processing in RELION. Acta Crystallogr D Struct Biol 73, 496–502 (2017).

66. S. H. W. Scheres, Amyloid structure determination in RELION-3.1. Acta Crystallogr D Struct Biol 76, 94–101 (2020).

67. S. Q. Zheng et al., MotionCor2: anisotropic correction of beam-induced motion for improved cryo-electron microscopy. Nat Methods 14, 331–332 (2017).

68. K. Zhang, Gctf: Real-time CTF determination and correction. J Struct Biol 193, 1–12 (2016).

69. T. Bepler et al., Positive-unlabeled convolutional neural networks for particle picking in cryo-electron micrographs. Nat Methods 16, 1153–1160 (2019).

70. K. Jamali, D. Kimanius, S. H. Scheres, in The Eleventh International Conference on Learning Representations. (2022).

71. P. Emsley, B. Lohkamp, W. G. Scott, K. Cowtan, Features and development of Coot. Acta Crystallogr D Biol Crystallogr 66, 486–501 (2010).

72. L. Hunter, T. E. Klein, in Biocomputing ’96. (WORLD SCIENTIFIC, 1995), pp. 1–774.

73. P. V. Afonine et al., Real-space refinement in PHENIX for cryo-EM and crystallography. Acta Crystallogr D Struct Biol 74, 531–544 (2018).

