## Supplemental Figures S1-S8, Tables S1-S5 for "Intact cells selectively amplify and structurally remodel amyloid fibrils"

**In cell NMR reveals cells selectively amplify and structurally remodel amyloid fibrils**

**The PDF file includes:**

Materials and Methods

Figs. S1 to S8

Tables S1 to S5

References 69-83

Materials and Methods

**Recombinant protein expression and purification**

Model building and refinement

For both Polymorph A and Polymorph B structures, an initial model was first built using Model Angelo (*80*) in the Relion pipeline, followed by manual building in Coot (*81*) and Chimera (*82*). The models were then refined with Phenix Real-space refinement (*83*). The statistics of the model validations are summarized in Table S2.

$y_{i}=A_{i} exp(-0.5(\frac{x-\mu_{i}}{\sigma_{i}})$^2^)

where $A_{i}$ is the amplitude, $\mu_{i}$ is the mean, and $\sigma_{i}$ is the standard deviation of the $i$-th Gaussian function. A constant offset $c$ was included to account for baseline correction. The total fitting function is given by:

$$y=y_{1}+y_{2}+y_{3}+y_{4}+c$$

An initial guess for the parameters was inputted, and the lower and upper bounds for the parameters were set to constrain the optimization process. The built-in MATLAB function ‘lsqcurvefit’ was used to fit the Gaussian functions to the data. The peak centers for the Cα and Cα’ peaks were fixed at 61.1 and 60.3 ppm, respectively, while the peak centers for the Cβ and Cβ’ peaks were allowed to vary.

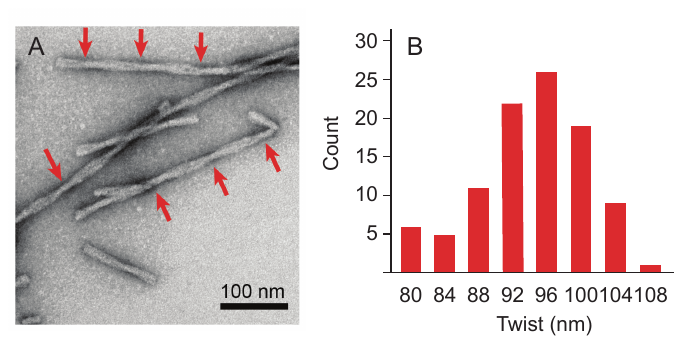

***Figure S1****: A) Negative stain transmission electron microscopy of Ac-α-syn(A53T) amyloid fibrils reveals a single major morphology with B) an average crossover length of 95 ± 2 nm.*

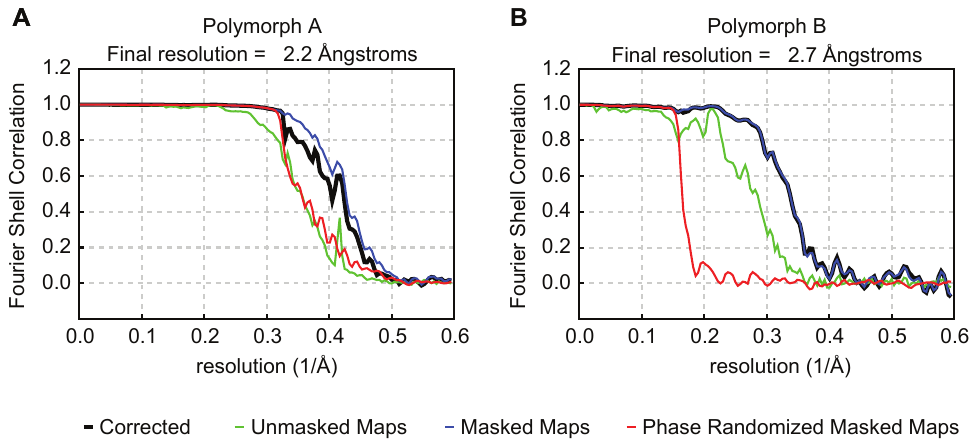

***Figure S2****: Fourier shell correlation curves for the cryo-EM structural refinement of A) polymorph A and B) polymorph B.*

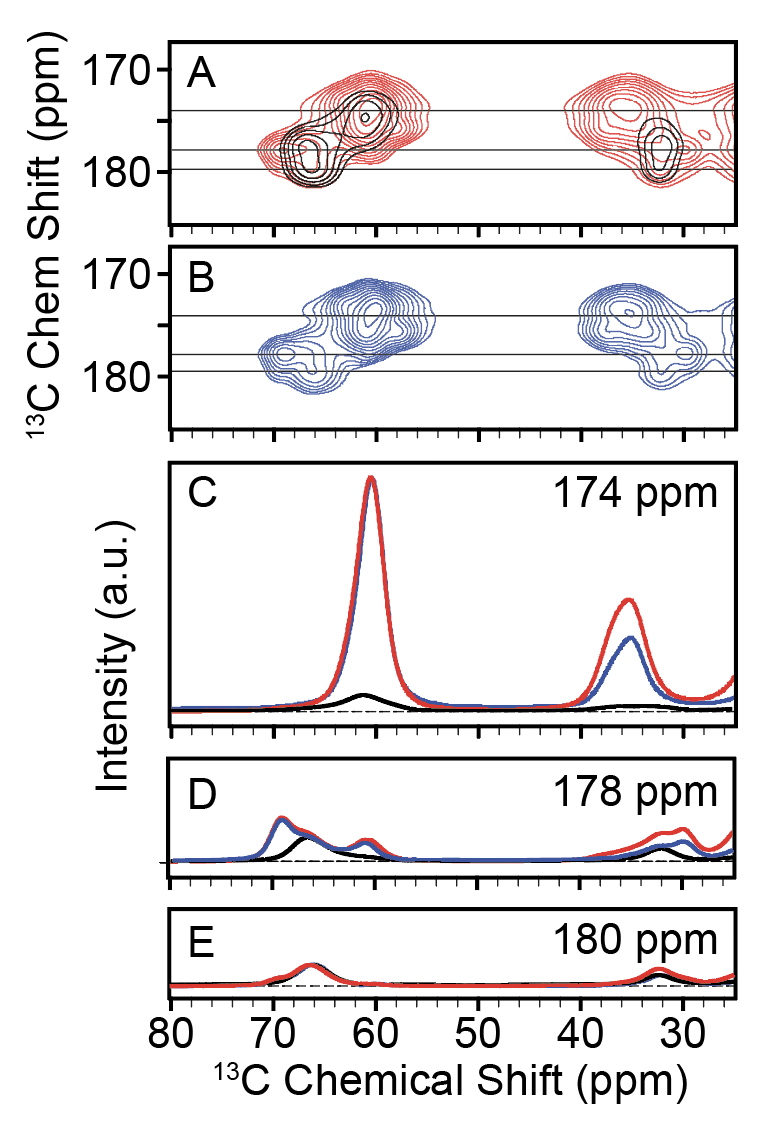

***Figure S3****: Ac-α-syn(A53T) fibrils seed true in vitro.* ***A****) Overlay of 2D ^13^C–^13^C of devo formed Ac-α-syn(A53T) fibrils seed that shows 19 valines (red) and only the four most N terminal valines (black), respectively. Average +/- two standard deviation are marked for each atom (black) as well as for the a-helical (red), random coil (green), and β (blue) secondary structures.* ***B****) 2D ^13^C-^13^C DARR spectra (20 ms mixing) of purified Ac-α-syn(A53T) monomers seeded into the fibril form with 1% (monomer/monomer) seeding of the de novo fibrils. One-dimensional slice from the 2D spectra at* ***C****) 174 ppm* ***D****) 178 ppm and* ***E****) 180 ppm, colored as in* ***A*** *and* ***B****.*

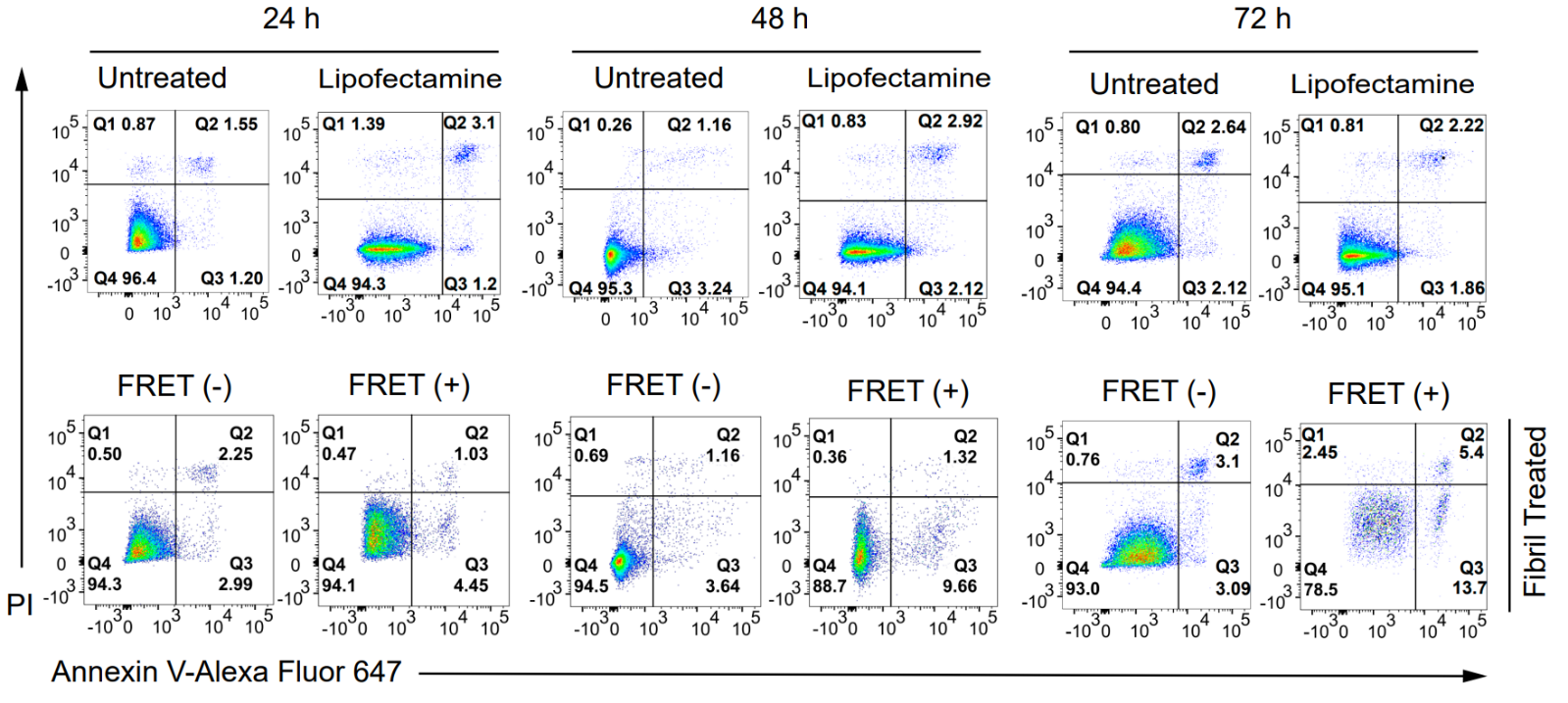

***Figure S4:*** *Primary FRET data to support results presented in Figure 3: α-syn aggregates increase cell death: Evaluation of apoptosis using Annexin V-PI double staining. HEK sensor cells were treated with 100 nM de novo fibrils and apoptosis was assessed by flow cytometry at 24, 48, and 72 h. The lower left and right panels represent FRET negative and FRET positive cell populations. The percentage of cells was characterized as those stained with both annexin-PI. The Q1, Q2, Q3, and Q4 quadrants represent necrotic, late apoptotic, early apoptotic, and live cell percentages, respectively. (*[*https://doi.org/10.1038/s41598-023-32075-9*](https://doi.org/10.1038/s41598-023-32075-9)*).*

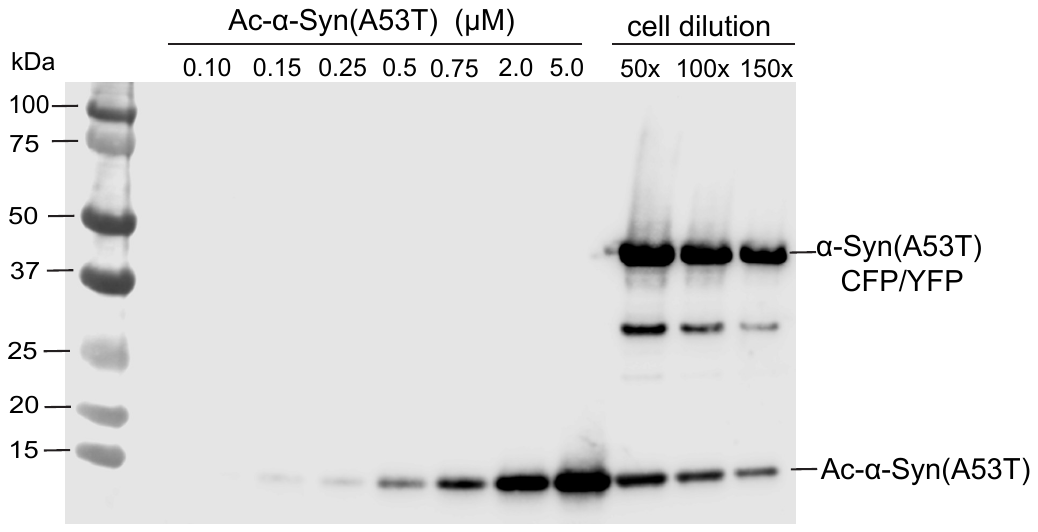

**Figure S5:** *Quantification of exogenously delivered a-syn(A53T) to HEK293T sensor cells by electroporation****.*** Western blot of lysed HEK293T biosensor cells that were electroporated with Ac-α-syn-A53T monomer. Intracellular protein concentration of Ac-α-syn(A53T) was determined by using the dilution series of lysed cells (lanes 10-12). Reference standard of recombinant Ac-α-syn(A53T) monomer was used for the calibration curve (lanes 3-9). Delivered intracellular Ac-α-syn(A53T) concentration was 70 ± 4 µM (n=3).

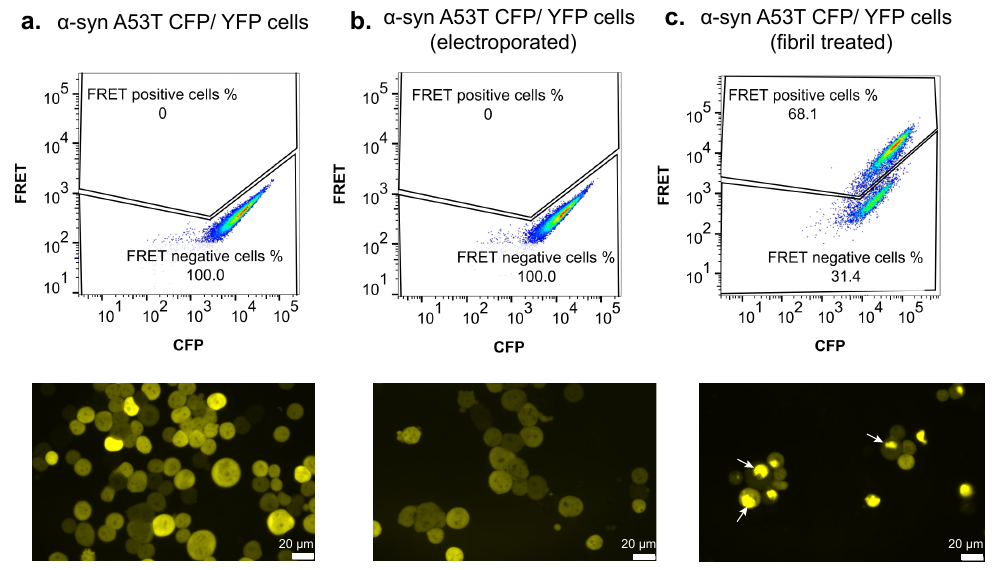

**Figure S6**: *No spontaneous FRET positive aggregate formation was detected by flow cytometry or microscopy after electroporation of HEK293 sensor cells that had not been exposed to fibrils with α-syn-(A53T) monomer.* ***A****) α-syn(A53T)-CFP/YFP HEK-293T biosensor cell line with no electroporation.* ***B****) α-syn(A53T)-CFP/YFP HEK-293T cell line electroporated in the presence of 1 mM with α-syn(A53T) monomer.* ***C****) α-syn(A53T) CFP/YFP HEK-293T biosensor cell treated with 100 nM sonicated Ac-α-syn(A53T) pre-formed fibrils. Representative images showing cells before electroporation (left), after electroporation in the presence of α-syn(A53T) monomer (middle), or after exposure to Ac-α-syn(A53T) fibrils (right). Arrows denote aggregates. Scale bars, 20 µm.*

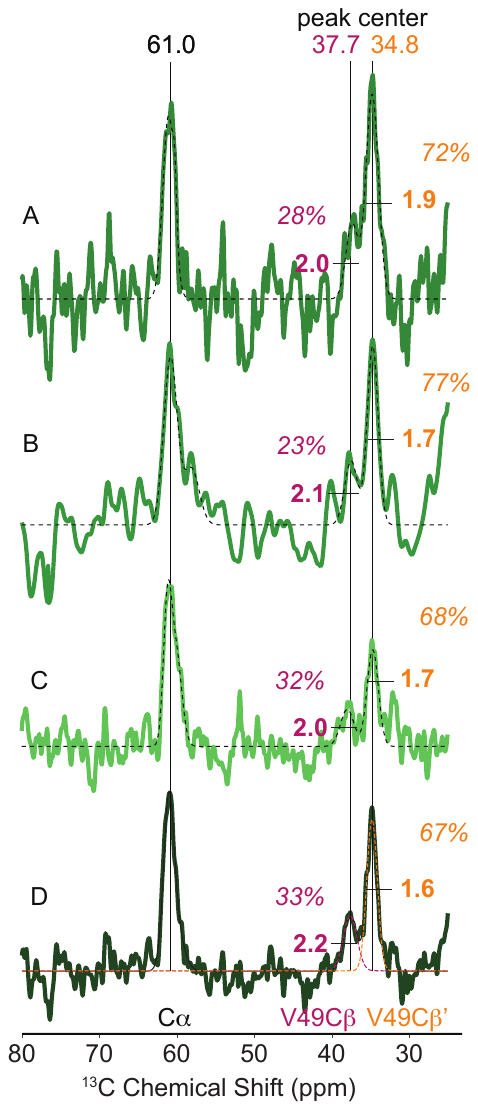

**Figure S7*:*** *1D NCOCx spectra that report on V49 in Ac-α-syn(A53T) fibrils that were propagated inside cells obtained on (****A-C****) three independent biological replicates and (****D****) the co-added spectrum of in cell propagated fibrils. The Cα peak was fit to a single Gaussian and the Cβ peaks were fit to two Gaussians. The FWHM are annotated in bold text and the percentage of the area of each Cβ peak are annotated in italics. Values V49Cβ are annotated in magenta and V49Cβ’ are annotated in orange. Fit parameters are reported in Table S4.*

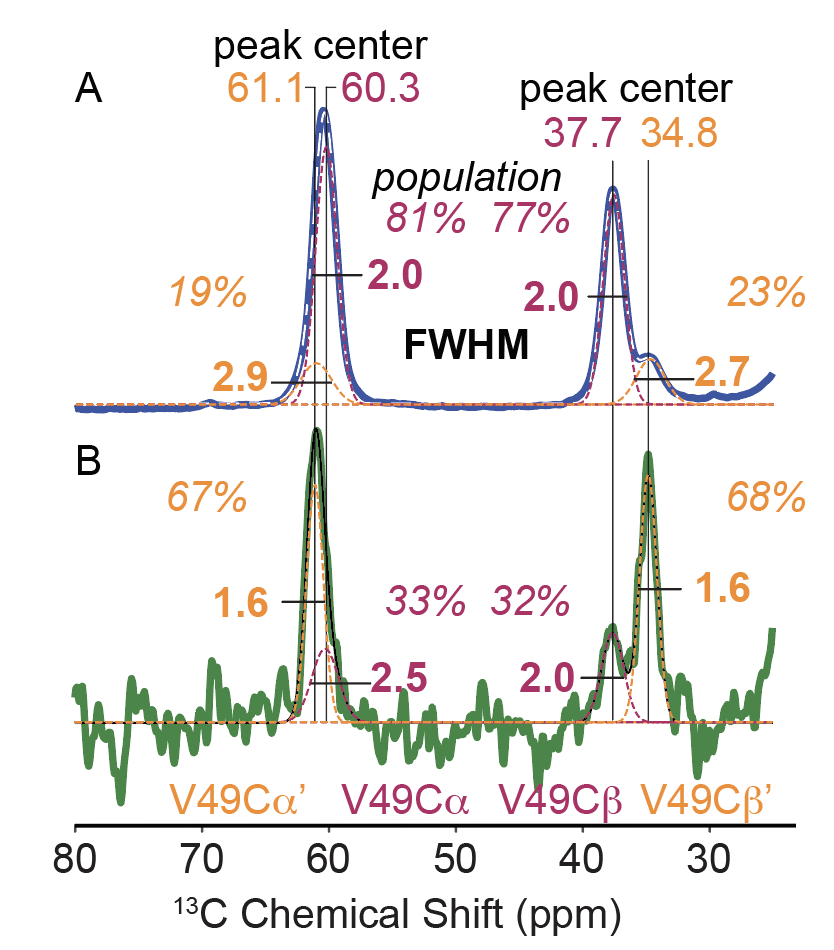

**Figure S8*:*** *1D NCOCx spectra that report on V49 in Ac-α-syn(A53T) fibrils that were propagated in Co-added spectra of* ***A****) de novo and* ***B****) in cell propagated fibrils with the Cα and Cβ peaks fit to a sum of two Gaussians. The centers for the Cα peak were fixed at 61.1 and 60.3 ppm. The other parameters were unconstrained. The FWHM are annotated in bold text and the percentage of the area for the Cα and the Cβ peaks are annotated in italics. Values V49Cα and V49Cβ are annotated in magenta and V49Cα’ and V49Cβ’ are annotated in orange. Values for the fits are also reported in Table S5.*

|  | Peak center (ppm) | | FWHM (ppm) | Population |
| --- | --- | --- | --- | --- |
| Cross-peak | C' | Cβ | direct dimension | (%) |
| V49C'-Cβ | 173.8 | 37.7 | 2.0 | 73 |
| V49C'-Cβ' | 174.0 | 34.9 | 3.1 | 27 |

**Table S1**: Chemical shifts of major and minor conformations of *de novo* fibrils as observed in the 2D NCOCx spectra.

| Microscope | Titan Krios (Thermo Fisher Scientific) | |
| --- | --- | --- |
| Camera | K3 (Gatan) | |
| Acceleration voltage (kV) | 300 | |
| Magnification | 105,000x | |
| Defocus range (µm) | -0.9 to -2.2 | |
| Dose rate (e^-^ /pixel/s) | 10 | |
| Number of movie frames | 50 | |
| Exposure time (s) | 3.5 | |
| Total electron dose (e^-^ /Å^2^) | 50 | |
| Pixel size (Å) | 0.84 | |
| Box size (pixel) | 300 | |
| Inter box distance (Å) | 38 | |
| Number of extracted segments | 2,266,580 | |
|  | **Polymorph A** | **Polymorph B** |
| Number of segments after 2D | 813,225 | 983,759 |
| Number of segments after 3D | 85,236 | 11,326 |
| Resolution, 0.143 FSC criterion (Å) | 2.21 | 2.68 |
| Map sharpening B-Factor (Å^2^) | -52.1 | -55.3 |
| Helical twist (°) | 179.61 | -1.21 |
| Helical rise (Å) | 2.40 | 4.81 |

**Table S2:** *Cryo-EM data collection and reconstruction parameters*

|  | **Polymorph A** | **Polymorph B** |
| --- | --- | --- |
| Non-hydrogen atoms | 4410 | 3750 |
| Number of protein chains | 10 | 10 |
| Map CC (mask) | 0.85 | 0.80 |
| RMSZ bonds (Å) | 0.005 | 0.005 |
| RMSZ angles (°) | 0.584 | 0.669 |
| All-atom clash score | 10.62 | 4.96 |
| Ramachandran favored/allowed/outliers (%) | 97.38/2.62/0 | 95.00/5.00/0 |
| Rotamer outliers (%) | 0 | 0 |
| Cβ outliers (%) | 0 | 0 |
| Molprobity score | 1.67 | 1.61 |

**Table S3:** *Cryo-EM modeling and refinement parameters*

| **Sample** | SNR | Peak Center (ppm) | | | | Fit | Full width half maximum (ppm) | | | | Population |
| --- | --- | --- | --- | --- | --- | --- | --- | --- | --- | --- | --- |
|  | at Cα | C' | Cα | Cβ | Cβ' | (*R^2^*) | C' | Cα | Cβ | Cβ' | of Cβ (%) |
| *de novo* fibrils in buffer | 260 | 174.1 | 60.4 | 37.6 | 34.6 | 0.98 | 3.5 | 2.2 | 2.2 | 2.4 | 78 |
|  | 279 | 174.0 | 60.4 | 37.6 | 34.7 | 0.98 | 3.5 | 2.2 | 2.0 | 2.9 | 79 |
| in buffer seeded fibrils | 233 | 174.2 | 60.3 | 37.7 | 34.4 | 0.98 | 3.7 | 2.1 | 2.1 | 2.4 | 81 |
|  | 185 | 174.3 | 60.3 | 37.6 | 34.2 | 0.97 | 4.0 | 2.4 | 2.4 | 2.4 | 82 |
| *de novo* fibrils diluted in lysates | 44 | 174.4 | 60.3 | 37.7 | 34.4 | 0.94 | 3.9 | 2.0 | 2.0 | 2.4 | 77 |
|  | 162 | 174.2 | 60.2 | 37.6 | 34.3 | 0.97 | 3.7 | 2.2 | 2.2 | 2.4 | 72 |
| in lysate seeded fibrils | 48 | 174.4 | 60.3 | 37.6 | 34.5 | 0.96 | 3.9 | 2.1 | 2.1 | 2.4 | 74 |
| in cell seeded fibrils | 7 | 175.4 | 61.0 | 37.5 | 34.8 | 0.59 | 6.0 | 2.0 | 2.0 | 1.9 | 28 |
|  | 7 | 176.3 | 60.7 | 37.5 | 34.8 | 0.76 | 8.0 | 2.3 | 2.1 | 1.7 | 23 |
|  | 10 | 176.0 | 60.8 | 38.0 | 34.7 | 0.75 | 6.5 | 2.1 | 2.0 | 1.7 | 32 |
| Co-addition of in cell seeded fibrils | 13 | 176.7 | 61.0 | 37.7 | 34.8 | 0.81 | 6.8 | 1.9 | 2.2 | 1.6 | 33 |

***Table S4****: Signal to noise ratio and results of fitting a 1D NCOCx spectra that report on V49 in Ac-α-syn(A53T) fibrils that were propagated in different environments to a signal Gaussian for the C’ and Cα peaks and two Gaussians for the Cβ peak.*

| **Sample** | SNR | Peak Center (ppm) | | | | Fit | Full width half maximum (ppm) | | | | Population | |
| --- | --- | --- | --- | --- | --- | --- | --- | --- | --- | --- | --- | --- |
|  | at Cα | Cα' | Cα | Cβ | Cβ' | (*R^2^*) | Cα' | Cα | Cβ | Cβ' | of Cα (%) | of Cβ (%) |
| *de novo* fibrils in buffer | 283 | **61.1** | **60.3** | 37.6 | 34.7 | 0.98 | 2.9 | 2.0 | 2.0 | 2.7 | 81 | 77 |
| in cell seeded fibrils | 13 | **61.1** | **60.3** | 37.7 | 34.8 | 0.81 | 1.5 | 2.9 | 2.2 | 1.6 | 38 | 33 |

***Table S5****: Signal to noise ratio and results of fitting co-added 1D NCOCx spectra that report on V49 in Ac-α-syn(A53T) fibrils that were propagated in different environments to two Gaussians with fixed centers (bold) for the Cα peak and two Gaussians for the Cβ peak.*
